# Inhibition of the SARS-CoV-2 helicase at single-nucleotide resolution

**DOI:** 10.1101/2022.10.07.511351

**Authors:** Sinduja K. Marx, Keith J. Mickolajczyk, Jonathan M. Craig, Christopher A. Thomas, Akira M. Pfeffer, Sarah J. Abell, Jessica D. Carrasco, Michaela C. Franzi, Jesse R. Huang, Hwanhee C. Kim, Henry D. Brinkerhoff, Tarun M. Kapoor, Jens H. Gundlach, Andrew H. Laszlo

**Affiliations:** Department of Physics, University of Washington, Seattle, WA 98195; Laboratory of Chemistry and Cell Biology, The Rockefeller University, New York, New York; Department of Biochemistry and Molecular Biology, Robert Wood Johnson Medical School, Rutgers University, Piscataway, NJ, USA

## Abstract

The genome of SARS-CoV-2 encodes for a helicase called nsp13 that is essential for viral replication and highly conserved across related viruses, making it an attractive antiviral target. Here we use nanopore tweezers, a high-resolution single-molecule technique, to gain detailed insight into how nsp13 turns ATP-hydrolysis into directed motion along nucleic acid strands. We measured nsp13 both as it translocates along single-stranded DNA or unwinds short DNA duplexes. Our data confirm that nsp13 uses the inchworm mechanism to move along the DNA in single-nucleotide steps, translocating at ~1000 nt/s or unwinding at ~100 bp/s. Nanopore tweezers’ high spatio-temporal resolution enables observation of the fundamental physical steps taken by nsp13 even as it translocates at speeds in excess of 1000 nucleotides per second enabling detailed kinetic analysis of nsp13 motion. As a proof-of-principle for inhibition studies, we observed nsp13’s motion in the presence of the ATPase inhibitor ATPγS. Our data reveals that ATPγS interferes with nsp13’s action by affecting several different kinetic processes. The dominant mechanism of inhibition differs depending on the application of assisting force. These advances demonstrate that nanopore tweezers are a powerful method for studying viral helicase mechanism and inhibition.

## Introduction

The severe acute respiratory syndrome coronavirus 2 (SARS-CoV-2) responsible for the COVID-19 pandemic has spurred large-scale efforts to develop antiviral drugs against its viral replication cycle (*1*). Once inside the host cell, the SARS-CoV-2 genome is translated by the host ribosome into two large polyproteins pp1a and pp1ab; the protease in pp1a, non-structural protein 5, (nsp5) then cleaves the polyproteins and releases additional nsps. Many of the nsps are critical for SARS-COV-2 replication, key among them are nsp5, the RNA dependent RNA polymerase: nsp12, and an RNA helicase: nsp13 (*2*). Antivirals (Paxlovid, Molnupiravir, and Remdesivir) currently approved to treat SARS-CoV-2 aim to inhibit viral replication by targeting nsp5 or nsp12, however, there are no presently approved antivirals which target nsp13. Studies of related coronaviruses (*3*) and other members of the order Nidovirales (*4*) have shown that the virus cannot replicate without a functional helicase. For this reason, nsp13 has been identified as a target for antiviral intervention. Moreover, unlike structural proteins, the amino acid sequence of nsp13 is one of the most conserved among closely related coronavirus species (e.g. SARS- and MERS-Nsp13) and newly emerging variants of SARS-CoV-2 (including Omicron (*5*)). Thus, nsp13 antivirals have the potential to be broad spectrum drugs that could address future coronaviral outbreaks as well.

Given this motivation, much has been learned about nsp13 in a relatively short time-span. Based on structure and biochemical studies, nsp13 has been classified as a SF1B-helicase and an inchworm mechanism for translocation has been proposed (*6–8*). In this mechanism, 2A and 1A domains each bind nucleic acids (NA) in a 5’ → 3’ orientation respectively (*6*). Binding of a nucleotide-triphosphate, e.g. ATP, is followed by 2A and 1A domains coming together in a “closed” conformation. During this conformational change, domain 2A advances a distance of one nucleotide in the 3’-direction along the NA strand. ATP hydrolysis then triggers a conformational change from “closed” to “open”, causing domain 1A to slide forward by a single nucleotide. This is followed by release of ADP and an inorganic phosphate to complete the cycle. Nsp13 can unwind both dsDNA and dsRNA in a 5’ → 3’ direction and is capable of hydrolyzing all deoxyribonucleotide triphosphates (dNTPs) and ribonucleotide triphosphates (NTPs) (*9–12*). Despite the RNA genome of SARS-CoV-2, structural data indicate that the nsp13 - nucleic-acid interactions are non-specific towards the ribose or nucleobase (*6*), and biochemical data show that the unwinding speed for dsDNA and dsRNA to be either similar or slightly faster for dsDNA (*9, 11, 12*). Reported unwinding speeds range from ~50-200 base pairs per second. On its own, nsp13 appears to be a relatively weak helicase with relatively low processivity during unwinding(*9, 12*). However, its processivity and unwinding speed are enhanced via destabilization of the DNA duplex using optical tweezers and interactions with RdRp (*7, 9, 12*). Translocation along ss NA is suspected since that is the mechanism by which nsp13 is thought to unwind NA duplexes but has not yet been observed. The speed at which nsp13 walks along its NA substrate in addition to its small step size puts the fundamental steps of the nsp13 helicase beyond the resolution of traditional singlemolecule techniques. Such resolution would enable detailed kinetic analysis of the underlying steps of nsp13’s inchworm mechanism and could also shed light on how inhibitory molecules affect its inner workings. We have developed a nanopore-based technique known as single-molecule picometer-resolution nanopore tweezers (SPRNT), which can measure steps of helicases and other molecular motors at high spatiotemporal resolution (*13, 14*). Here we apply SPRNT to nsp13 of SARS-CoV-2 to reveal the detailed mechanism of nsp13’s motion on DNA and we demonstrate how SPRNT can be used to determine the mechanism of action of a helicase inhibitor.

In SPRNT, a single *Mycobacterium smegmatis* porin A (MspA) nanopore within a phospholipid bilayer serves as the sole electrical connection between two electrolyte-containing wells (Fig. 1A). A voltage applied across this membrane causes an ion current to flow through the nanopore. The electric field of the nanopore also draws negatively-charged NA through the pore, blocking the ion current. Different NA bases within the nanopore’s constriction cause unique ion current blockages which can be decoded into NA sequence as in nanopore sequencing (*15*). If a helicase molecule is bound to the captured NA strand, the helicase will come to rest on the outer-edge of the entry into the nanopore (henceforth, referred to as pore-rim) as the helicase’s large size prohibits entry into the pore. From this point onward, the helicase pulls the NA through the nanopore against the electrostatic force on the NA, resulting in a sequence of ion current steps as the NA sequence in the pore changes. These ion current steps resolve single-nucleotide steps on sub-millisecond time scales, making it possible to observe the details of helicase motion along NA, while simultaneously determining the NA sequence of the substrate in the helicase. The force SPRNT applies on the enzyme/NA complex is proportional to the applied voltage. Depending on which end of the NA is threaded into the pore, this force can either oppose or assist the helicase motion.

**Figure 1:**
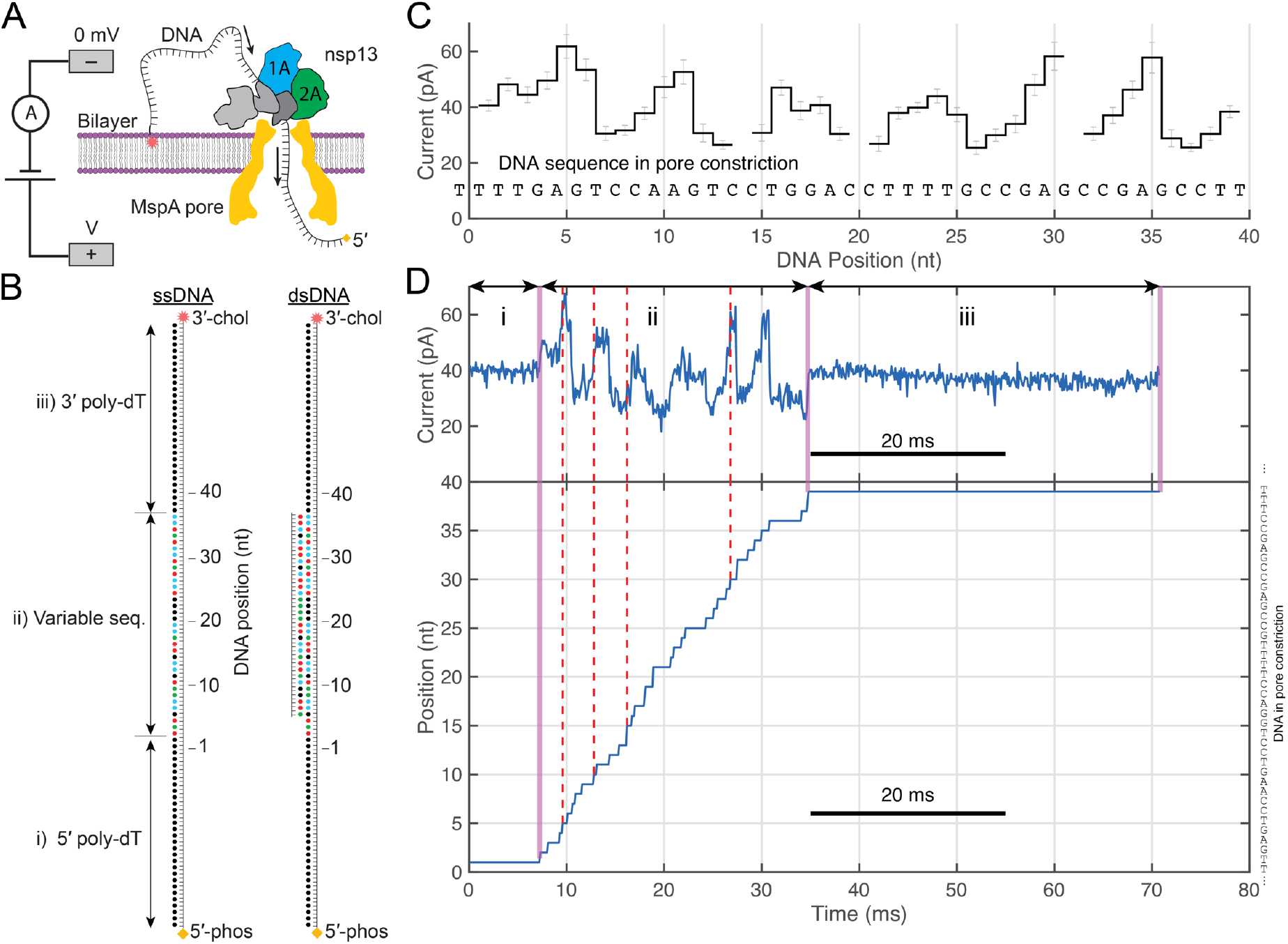
Nanopore tweezers with nsp13 helicase. A) Schematic of the nanopore experiment. B) Schematic of the DNA used in this manuscript. The template DNA strand has both 5’- and 3’-dT30 tails with a region of 36 nucleotides of variable sequence in the middle. dsDNA experiments used a 32 nucleotide complementary strand. A 3’-cholesterol causes the DNA strand to associate with the bilayer increasing the local concentration of DNA near the nanopore in order to increase the rate of DNA capture. While the additional negative charge of a 5’-phosphate helps to encourage feeding of the 5’ end of the DNA into the nanopore. As an nsp13 molecule feeds a DNA strand into the pore ion-current levels corresponding to i) the 5’-dT30, ii) the variable sequence region, and iii) the 3’-dτ3θ are observed in the ion-current measurement. C) Ion-current consensus for the DNA sequence shown as it translocates through the nanopore constriction. For the shown section of 40 DNA bases, 37 ion current levels are regularly found within the ion current traces. This occurs when currents for adjacent k-mers are indistinguishable. Gaps in the consensus represent these unobserved steps. D) Example ion current trace for nsp13 walking along the ssDNA strand. i) the 5’ poly-dT section is within the pore, ii) the variable current section is within the pore, iii) the 3’ poly-dT section is within the pore. The variable current region of the trace can be aligned to the ion current consensus in C) to produce what is essentially a best fit for the enzyme’s location along the DNA strand as a function of time (below). Red dashed lines are guides to the eye showing where the 5^th^, 10^th^, 15^th^, and 30^th^ levels appear within the read as determined by the automatic alignment.

## Results and Discussion

Because nsp13 is known to unwind DNA(*9, 12*), has a higher affinity for DNA(*9*), and working with DNA is easier and more affordable, we designed a DNA substrate containing a 37-nucleotide pseudorandom sequence flanked by 30-nt poly-T overhangs on the 3’- and 5’ ends (Fig. 1B). A cholesterol tag at the 3’ end of the DNA strand was used to increase the DNA concentration near the bilayer, thereby increasing the rate of capture into the pore. The DNA base sequence of the pseudorandom section was chosen to have varied sequence context and easily distinguishable ion current levels (Fig. S1). A 32-nucleotide complementary strand could be hybridized to this strand for dsDNA experiments (Fig. 1B, Table S1, Fig. S2, Materials and Methods). Nsp13 loads onto the 5’ tail near the dsDNA-ssDNA junction and the free 5’ end of the DNA then threads into the nanopore. We recorded a total of 2413 individual nsp13 translocation and unwinding events and a total of 27,641 helicase steps (Table S2). Both ssDNA and dsDNA templates reveal 1-nt steps in a 5’ → 3’ direction with significantly slower nsp13 motion on dsDNA (Figs. 1C, 1D, S1, S2). At saturating [ATP] and ~36 pN assisting force, we observe nsp13 ssDNA translocation speed of 1140 ± 40 nt/s (mean ± s.e.m) and an average dsDNA unwinding speed of 120 ± 10 bp/s (mean ± s.e.m) (Fig. 1E & Fig. S4). In both translocation and unwinding experiments, a broad range of speeds is observed between individual helicases, suggesting a significant amount of static disorder affecting the helicases’ function. Even during the fast nsp13 translocation of 1500 nt/s, single-nucleotide steps are well resolved (Fig. 1D).

The remarkable speed of nsp13 during translocation presented an analysis challenge as steps within the data were at times difficult to resolve. In the past, our analysis of nanopore tweezers data involved first automated level-finding using a change-point algorithm followed by alignment of the found levels to the ion-current consensus (*13, 14, 16*). In that analysis scheme, level-finding was agnostic to the underlying DNA sequence and detection of levels was achieved by looking for statistically significant steps within the data (*17*). Here, nsp13’s speed makes such step detection difficult or impossible as individual steps sometimes consist of just a few data points which is insufficient to agnostically determine whether a step took place. However, here the DNA sequence, and therefore the expected progression of ion current levels, is known (Fig. 1D) and that information can be used to inform step-finding. By adapting our alignment algorithm so that raw data was aligned to the ion-current consensus, we were able to perform sequence-aware step finding (Figs. S5, S6, S7). In this scheme, the ion-current datapoints for a given read are aligned to consensus current values. The resulting alignment (Fig. 1D, *bottom*) yields what is essentially a global best-fit location of the nsp13 helicase as a function of time. We can then extract kinetic parameters such as step dwell-time, the location and frequency of enzyme backsteps, and the location of read start and read terminations. Such automated alignments can be readily verified by-eye, Supplemental Figures S6 and S7 show several representative alignments for ssDNA translocation and dsDNA unwinding, respectively.

### Single-nucleotide ssDNA translocation

Figure 2A shows the dwell-time distribution for 4 representative locations along the DNA strand. Intriguingly, nsp13 steps over each of these locations at different rates. Such sequence-dependence has been reported for other helicases (*14, 18–20*). At saturating conditions, the underlying dwell-time distributions of each nucleotide transition are well-fit by a single exponential function (Fig. 2A, Fig. S8), indicating that each step during nsp13 translocation is governed by a single rate-limiting step. Since processive helicases couple ATP hydrolysis to DNA translocation, we varied the ATP concentration to probe nsp13 translocation kinetics. The rate of translocation is dependent on [ATP] and is well described by Michaelis–Menten kinetics, with the maximum reaction rate (Vmax) ranging from 600 to 3000 s^−1^ and with Michaelis constant (Km) ranging from 100 to 700 μM for ATP (Fig. 2B, Fig. S9) depending on the underlying sequence context within nsp13. The large variation in rate of translocation at different DNA positions suggests that the nucleic-acid base identity affects nsp13 translocation kinetics.

**Figure 2:**
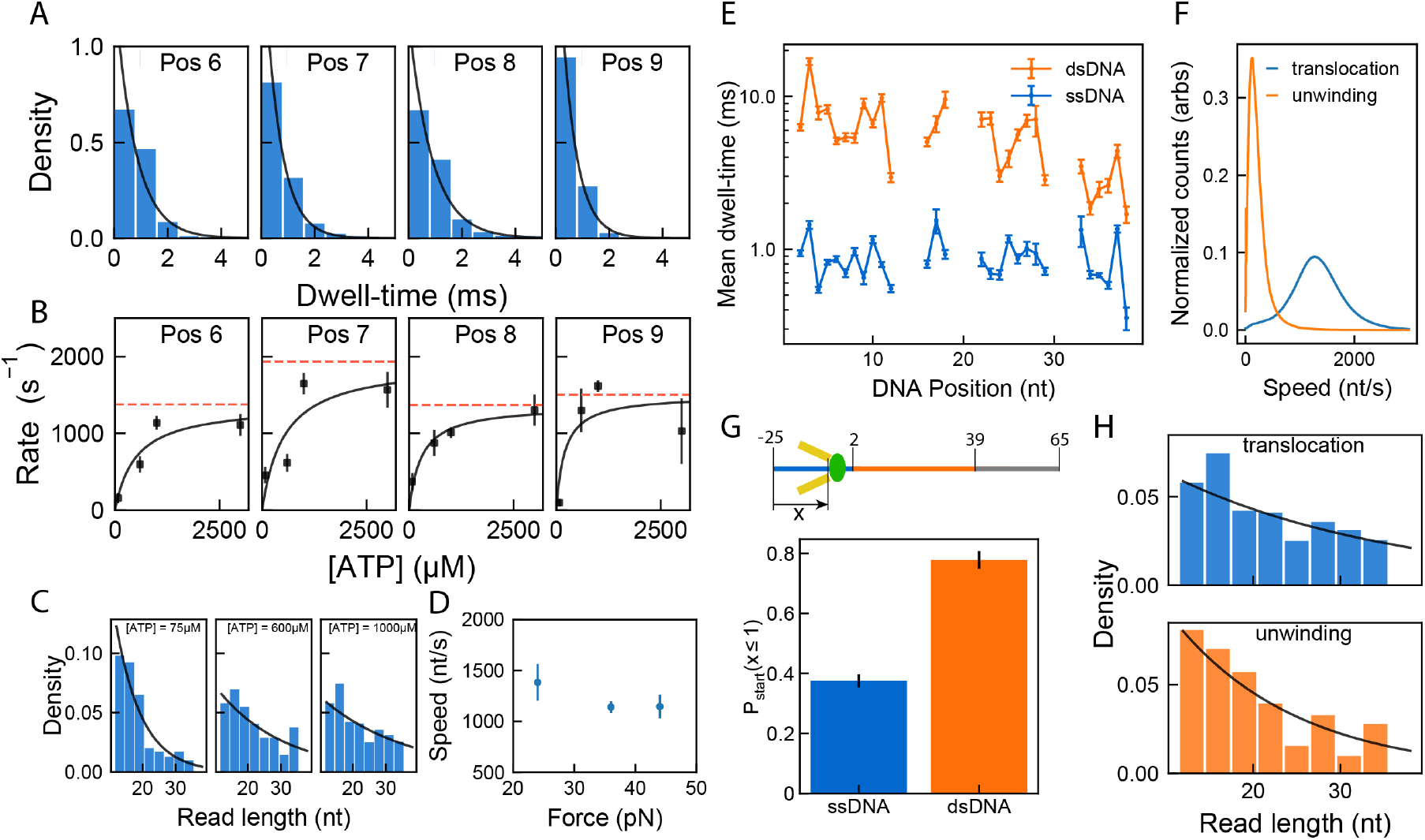
Sequence-dependent nsp13 translocation and unwinding kinetics. A) Distributions of dwell-time for a few positions from all translocation events at 1000 μM ATP. An exponential fit to the data is shown in black. B) Enzyme rate as a function of [ATP] at corresponding DNA positions from (A). Rate is calculated as the reciprocal of mean dwell-time at the DNA position. The mean dwell-time at each [ATP] was calculated by taking the average of all dwell-times from t=0 s to t=100 s. The black line shows the best fit of the data to the Michaelis-Menten equation. The red line corresponds to maximum velocity of reaction (Vmax). C) Comparing processivity (defined as bases translocated before falling off) for three different ATP concentrations: 75 μM, 600 μM and 1000 μM ATP (n=384, 162 and 825 events respectively). The black line shows the best fit of the data to the processivity equation defined in Supp. Disc. X. D) The effect of applied force on nsp13 translocation on ssDNA was studied by varying the voltage at constant saturating [ATP]=3 mM (A) Mean dwell-time (ms) (mean ± s.e.m) as a function of force (pN) measured at 3 different voltages 220 mV, 180 mV, 120 mV that corresponds to ~44 pN, ~36 pN, ~24 pN force respectively (n=1794, n=4329, and n=507 events respectively). E) Mean dwell-time of nsp13 at each DNA position measured in Figure 1. for translocation (blue) and unwinding (orange), at 1000 μM ATP (n=819 and 902 events respectively). Data are means ± SEM. Gaps are DNA positions where the measured ion-current between two nucleotide positions was identical and a mechanical step could not confidently be identified. F) Comparing speed of translocation vs. unwinding. G) Probability of starting at DNA position <= 1 for translocation and unwinding. (Above) Schematic showing DNA positions, numbered with respect to start and end of double-stranded unwinding region. H) Probability distribution of the length along the DNA over which the nsp13 stayed bound (“read length”, processivity) for translocation vs. unwinding.

Nsp13 translocation can begin and end anywhere on the template DNA, with 50% of reads terminating after nsp13 walkings into the 3’ poly-dT section at the end of the DNA template (Fig. S10). The true processivity of nsp13, defined as the number of bases translocated before nsp13 disassociates, is estimated by fitting the observed read-lengths at varying [ATP] (Supp. text. and Fig. 2C). Processivity at saturating [ATP] is 26 ± 4 nt and drops to 7.9 ± 0.7 nt (mean ± stdev) at limiting [ATP] = 75μM. While nsp13 is shown to bind DNA in the absence of ATP (*9*), our data suggests that while waiting for ATP to bind, the nsp13-DNA complex is less stable and more easily dislodged by application of force.

During SPRNT, the electrostatic force pulling on the NA by the electric field in the pore constriction leads to a contact force on the surface of the enzyme by the pore-rim. In our experimental design, this pushing force on the helicase (~36 pN at 180 mV) acts on domain 2A (Fig. 1A & Fig. S3). With the 5’ end of the NA in the pore, the force on the NA acts in the direction of motion of the enzyme. Previous nanopore tweezers experiments have seen evidence for a one-dimensional diffusive step in which the applied force, rather than ATP-hydrolysis drives enzyme motion. To investigate the affect of assisting force on nsp13 translocation, we varied the force from ~24 pN to ~44 pN at saturating [ATP] and found no significant change in average speed (Fig. 2D and Fig. S11, S12). Since the speed of translocation is independent in the range of forces we studied but dependent on [ATP], these data suggest that the observed nsp13 translocation is predominantly on-pathway ATP-hydrolysis-driven motion.

By definition, at saturating [ATP], ATP-binding is not rate-limiting. This, in combination with the lack of apparent force dependence, suggests that either (i) ATP-hydrolysis, (ii) domain 1A motion, or (iii) ADP & phosphate release are rate-limiting for nsp13 translocation. The apparent sequence dependence of this rate-limiting step suggests that it is domain 1A motion, but these observations do not strictly rule out effects of the DNA sequence on the rate of ATP hydrolysis or ADP release.

To further investigate the ADP-release portion of nsp13’s kinetic pathway we also ran the ssDNA assay in the presence of equimolar ATP and ADP ([ATP] = [ADP] = 600μM). In these conditions, the rate of nsp13 translocation was slowed by approximately a factor of two (Fig. S13). This is similar to nanopore tweezers results with Hel308 helicase and PcrA helicase where ADP was seen to act as a competitive inhibitor to ATP-binding(*14, 18*). Similar to force-assisting nanopore tweezers experiments with PcrA, ADP causes an increase in step dwell-time. ADP-binding does not appear to be associated with a physical step of the DNA through the pore.

### Double stranded DNA unwinding

We next observed nsp13’s ability to unwind dsDNA. The mean dwell-times for duplex unwinding steps are similarly sequence-dependent and on average ~8-fold longer than that of ssDNA translocation (Fig. 2 E, F).

While ssDNA reads start at a broad range of DNA positions, nearly 80% of the dsDNA reads start with the 5’ poly-dT within the pore constriction (Fig. 2G and Fig. S14). The reads that start within the duplex region are also comparable in speed to dsDNA unwinding (Fig. S15). This suggests the majority of the reads, including the partial dsDNA reads, are unwinding reads and that limited nsp13 unwinding occurs in the bulk solution.

Compared to ssDNA translocation, the processivity during dsDNA unwinding is shorter at 14 ± 1 nt (mean ± stdev) (Fig. 2H), which is consistent with previous results showing low processivity during unwinding (*9*). The underlying dwell-time distributions are single-exponential (Fig. S16), indicating a single, rate-limiting step. Because dsDNA unwinding is so much slower than ssDNA translocation, it stands to reason that the rate-limiting substep is the sub-step in which DNA unwinding occurs. Intriguingly, the dwell-times for dsDNA unwinding and ssDNA translocation as a function of DNA position are correlated (Fig. 2E). A similar effect was recently reported with the superfamily 1A helicase PcrA (*14*). In that case, it was deduced that such a correlation would occur if unwinding and translocation shared the same rate-limiting step and the presence of the DNA duplex acts as an additional barrier to that rate-limiting step. The duplex then modifies the rate-constants for the rate-limiting step rather than adding a new rate-limiting step. From this we can conclude that the advance of domain 1A following ATP-hydrolysis is rate-limiting in both translocation and unwinding and that unwinding occurs coincident with this step.

Based on these data and in light of insights from studying other SF1 and SF2 helicases using SPRNT (*13, 14*), we propose an ATP hydrolysis model that places the steps of the inchworm mechanism in the context of the observed transitions during SPRNT (Fig. 3). In SPRNT, only steps in which DNA moves relative to the pore produce observable steps in the data. In the experiment shown here, domain 2A rests on the pore-rim, meaning that only the motion of DNA relative to domain 2A is observable (*14*). This is also the only kinetic step that can be modified by the applied force (*21*). Other kinetic sub-steps, along the rows of Fig. 3, are inferred from the underlying dwell-time distributions and the known steps of the inchworm mechanism (*18*). For example, the presence of equimolar ADP slows nsp13 by encouraging binding of ADP, thereby backtracking nsp13 on its kinetic pathway from state I to state V. This does not result in DNA motion through the pore but does increase the step dwell-time.

**Figure 3:**
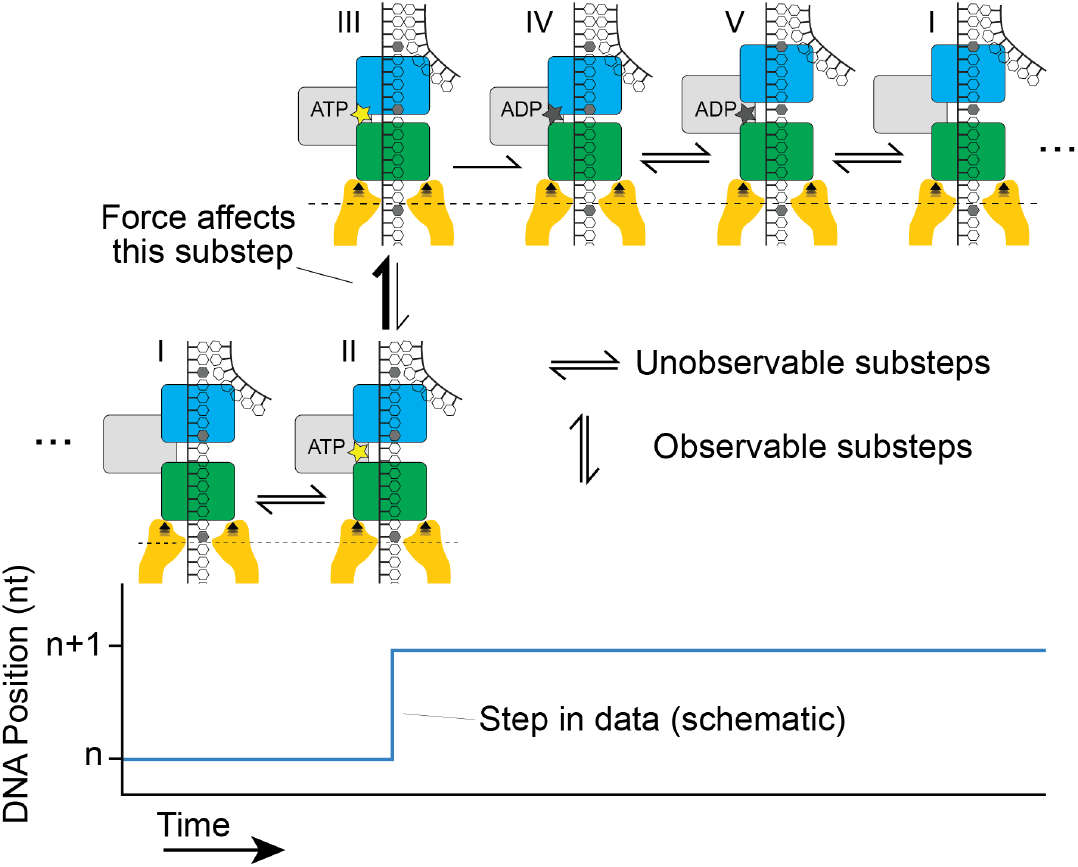
Nsp13 ATP-hydrolysis mechanism in the context of SPRNT. The observable chemical step of the ATP hydrolysis cycle during a SPRNT measurement is on the y-axis. Inferred steps, not associated with mechanical DNA motion, are on the x-axis. Chemical steps based on inchworm mechanism are: I) Enzyme bound to DNA, walker domains apart (green: domain 2A, blue: domain 1A); II) Nucleotide triphosphate bound, walker domains apart; III) Nucleotide triphosphate bound, walker domains together; IV) ADP bound, walker domains together; V) ADP bound, walker domains apart; the assisting force of SPRNT can only affect steps in which DNA moves through the pore; i.e. transitions between states II and III.

### Nsp13 inhibition by ATPγS

There is significant interest in determining mechanisms by which nsp13 can be inhibited by small molecules (*12, 22*). Because SPRNT can resolve individual ATPase turnovers, it can provide detailed insight into the mechanism of action of inhibitory molecules. As a proof of principle, we investigated inhibition of nsp13 by using the slowly-hydrolysable ATP-analog ATPγS (*23, 24*). With ATPγS present, the average speed and processivity of nsp13 is significantly reduced relative to the ATP-only condition for translocation (Fig. 4A-E). Commercially available ATPγS regularly contains ADP as a significant contaminant comprising as much as 10% of the reagent, however control experiments with a mixture of 600μM ATP and 600μM ADP showed significantly less inhibition (Fig. S13B), suggesting that the observed inhibition was indeed due to ATPγS. To determine whether ATPγS alone was sufficient for ssDNA translocation, we attempted an experiment with [ATPγS] = 600 μM and observed translocation with similar speed and processivity to ATPγS and ATP mixed condition in which [ATP] was 2x higher than [ATPγS] (Fig. 4A,B,E). This suggests that nsp13 is capable of hydrolyzing ATPγS and that the inhibitory effect of ATPγS saturates above 300μM suggesting that nsp13 has a higher affinity for ATPγS than for ATP. Dwell-time distributions for nsp13 translocation on ssDNA with ATPγS are well described by an exponential distribution suggesting the presence of a single rate-limiting step (Fig. S17). Intriguingly, the rate constants for each DNA position are different, suggesting that the DNA sequence within the nsp13 affects this rate-limiting step.

**Figure 4:**
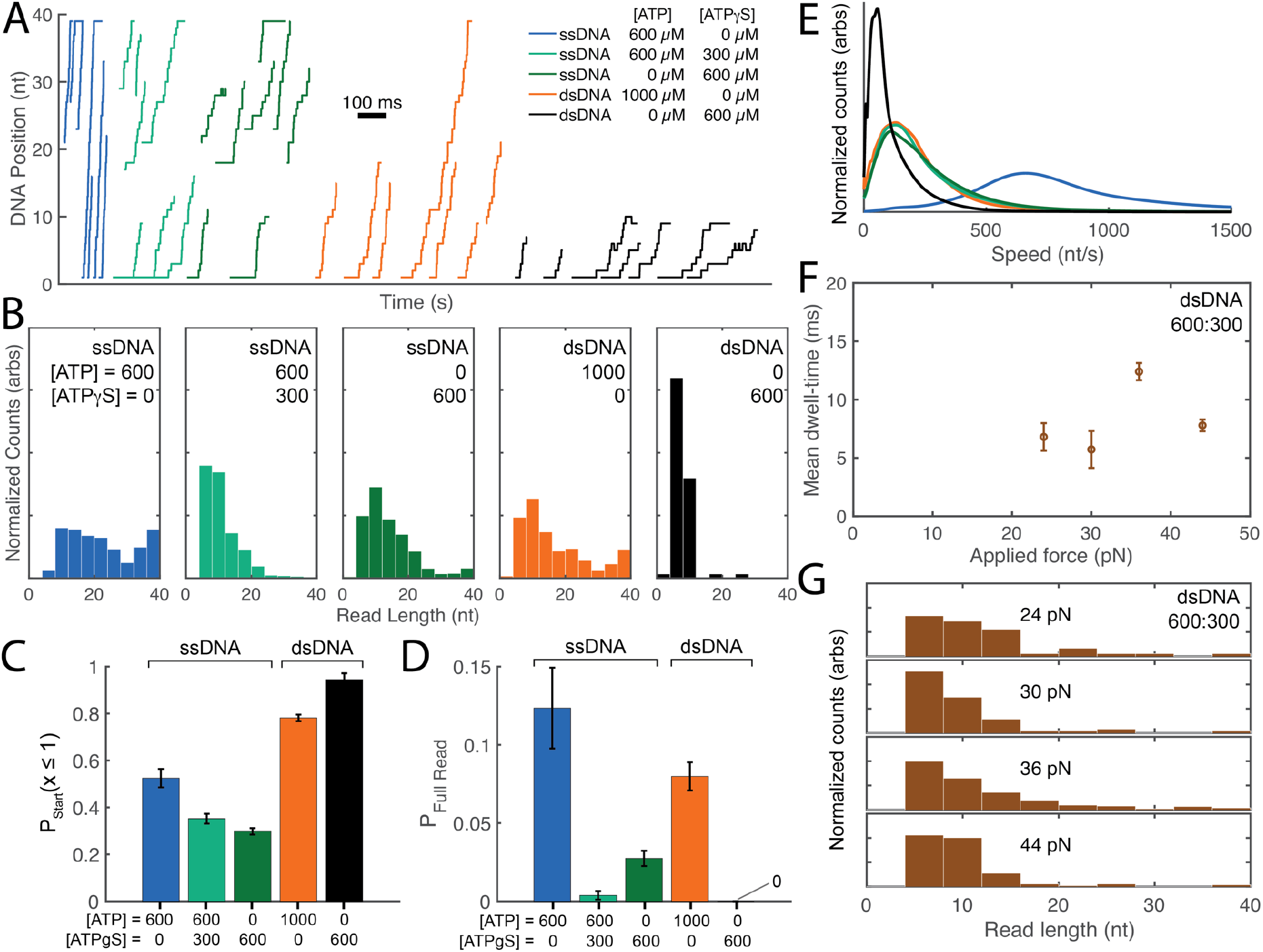
ATPγS inhibition of nsp13. A) Example position versus time traces for nsp13 unwinding events in the presence of ATP only (blue), ATP and ATPγS (red) and ATPγS only (black). B) Distribution of read-lengths (each a measure of the distance enzyme travelled before disassociation) during translocation and unwinding for ATP and ATPγS mixed conditions. C) Probability of an enzyme event starting at DNA position ≤ 1 for corresponding events in (B). D) Probability of a full read for corresponding conditions in (C). E) Comparison of speed of translocation and unwinding during translocation and unwinding in various mixed ATP and ATPγS conditions.The effect of applied force on nsp13 unwinding activity was measured in the presence of [ATP] = 600 μM and [ATPγS] = 300 μM. F) Mean dwell time per event averaged over all DNA positions at various assisting forces. All error bars are SEM. G) Distribution of the measured read-lengths for the corresponding nsp13 events at the indicated forces.

We also tested nsp13’s ability to unwind the DNA duplex in the presence of ATPγS. Unwinding speed and processivity are significantly reduced by the presence of ATPγS. In ATPγS-only conditions, nearly 100% of unwinding reads begin with the 5’ poly-dT within the nanopore constriction, (significantly greater than was observed in conditions that included ATP) indicating that without ATP present, there is no significant unwinding in bulk. This suggests that nsp13 can only unwind DNA using ATPγS in the presence of an assisting force provided by the nanopore.

We further investigated the role of this assisting force for ATPγS-associated motion by varying force from 24 pN to 44 pN at constant [ATP] (600 μM) and [ATPγS] (300 μM). At all applied forces, there was no significant change in average speed (Fig. 4F) or processivity (Fig. 4D). As noted above, we expect the force exerted in this experiment to affect only the kinetic step in which domain 2A advances along the DNA following NTP binding. In sum, we make the following observations: 1) ATPγS slows the progression of nsp13 on ssDNA and dsDNA, 2) inhibition by ATPγS saturates above ~300 μM, 3) there is little force dependence in the range 24 pN – 44 pN, and 4) with zero assisting force (bulk) ATPγS is insufficient for unwinding while with assisting force (on the nanopore) nsp13 can use ATPγS to unwind DNA.

Using these observations, we have constructed a model of ATPγS inhibition for nsp13 (Fig. 5). ATPγS inhibits nsp13 unwinding in three ways. First, ATPγS reduces nsp13’s processivity. Second, at zero force, ATPγS inhibits the first conformational change/step of the inchworm mechanism. This could be either due to reducing the closing-rate of domain 1A and 2A to essentially zero or due to destabilizing the closed state so that it rapidly backsteps to State II before ATPγS can be hydrolyzed. Third, even in the presence of an assisting force, which helps to stabilize nsp13 in the closed conformation (State III), ATPγS-hydrolysis is slower than ATP-hydrolysis. We hypothesize that this is the principal reason why ATPγS slows nsp13 in the SPRNT assay. Alternative models in which ATPγS is not hydrolyzed are described in Fig. S18, however only the model presented in Fig. 5 is capable of describing all of our observations (Figs. S18 and S19).

**Figure 5:**
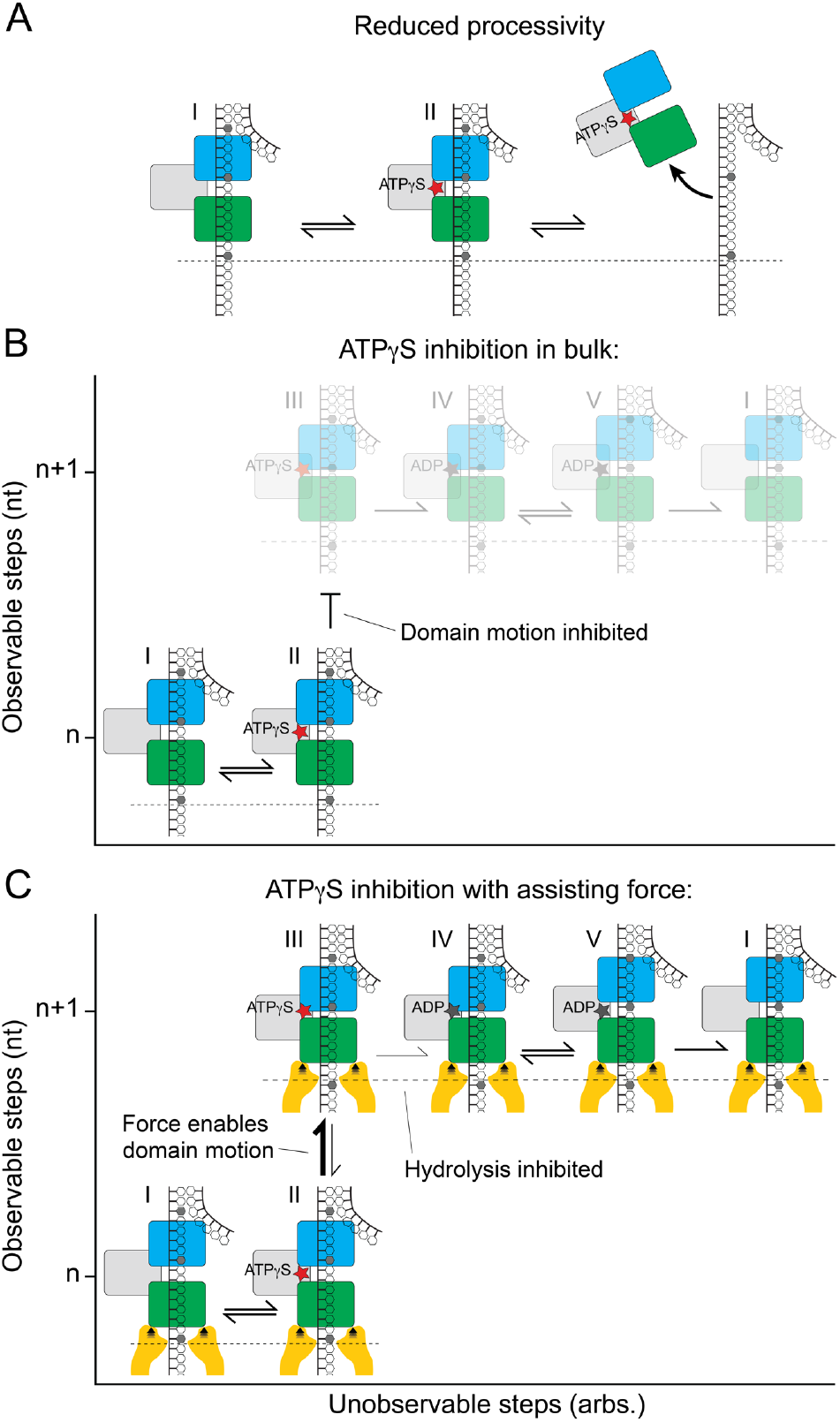
Model of nsp13 ATPγS inhibition. ATPγS inhibits nsp13 via three separate mechanism. A) nsp13 has lower processivity with ATPγS. B) Without assisting force (in bulk), ATPγS inhibits the motion of domain 2A following ATPγS-binding. C) The assisting force applied in SPRNT overcomes the inhibitory effect of ATPγS seen in B), however, hydrolysis of ATPγS is also slower as is seen in many other ATPases.

This model can also be used to make predictions for the combined effects of ATPγS and the presence of a DNA duplex on the rate of nsp13 progression. In other words we can use observations of dsDNA unwinding (Fig. 4 *orange*, Fig. S16) and ATPγS-driven ssDNA translocation (Fig. 4 *dark green*, Fig. S17) to predict nsp13’s behavior while unwinding with ATPγS-alone (Fig. 4 *black*). In this model, ATPγS-hydrolysis and duplex unwinding are sequential steps and are a similar order of magnitude. Therefore, the mean step dwell-time for dsDNA unwinding with ATPγS-alone is given by the sum of the mean step dwell-time for unwinding and the mean step dwell-time for ATPγS-driven ssDNA translocation and the dwell-time histograms at each position should take the form of a convolution of the dwell-time histograms from Figures S16 and S17. The data are in good agreement with this prediction (Fig. S20).

## Discussion

While previous biochemical data show nsp13 ATPase rate is enhanced up to 25-fold in the presence of singlestranded nucleic acid (*10*), there has been no measurement of nsp13 translocation speed until now. Unwinding speed is ~8-fold slower compared to translocation which is similar to that recently reported for SF1A helicase PcrA using SPRNT (~4-fold), and consistent with *in vitro* studies of other passive helicases (*14, 25, 26*). Notably, the nsp13 unwinding speed measured here is similar to the observed speed of SARS-CoV-2 RdRp with which nsp13 forms a complex: ~170 nt/s at 37 °C as determined by single-molecule magnetic tweezers (*27*).

Our assisting-force unwinding data shows speeds about 10-fold greater than estimates of bulk unwinding speed. However, our average unwinding speed is comparable to the unwinding speeds achieved using optical tweezers to destabilize the RNA duplex (*9*). The manner by which force is applied with SPRNT differs significantly from how force was applied in the optical tweezers study (Fig. S21). In SPRNT, force affects the advance of domain 2A following ATP-binding whereas in the optical tweezers assay, force destabilizes the RNA duplex. It is possible that in the absence of force, domain 2A advance is rate-limiting as is apparent with ATPγS at zero force (bulk). Still, all other kinetic steps observed here are unaffected by the force applied in the SPRNT experiment, and even the application of an assisting force to nsp13 may have biological relevance. Interactions between nsp13 and the SARS-CoV-2 RdRp have been shown to play regulatory roles that affect unwinding activity (*6, 7, 12, 28, 29*). Cryo-EM structures with nsp7, nsp8, and nsp7 show a pair of nsp13 associated with the RTC in which one nsp13 is also bound to an RNA strand. In these structures, the RNA-bound nsp13 has a 1B-domain which is swiveled closer to domains 1A-2A while in the other the 1B domain is further apart from domains 1A-2A. These states are stabilized sterically by the second nsp13 molecule in the RTC. This suggests that interaction with the 1B-domain, such that it is in a closed state, can favor RNA binding and conceivably lead to a more processive helicase (*6*). Similarly, engineering the DNA groove to be in a closed conformation has been shown to increase unwinding activity for SF1A helicases such as PcrA and Rep (*30*). Contact between nsp13 and the nanopore rim may be playing a similar role. A similar study in which the conformation of the nsp13 1B-domain is constrained could shed further light on the role of the 1B-domain.

Unlike many single molecule techniques, SPRNT is uniquely capable of studying kinetics of translocation of a helicase in the absence of a duplex. While in this study, nsp13 translocation was measured only with assisting forces, reversing the orientation such that nsp13 is pulling DNA out of–rather than lowering it into–the pore would create an opposing force on domain 1A (*14*). This would directly mimic how the RTC complexes may apply force *in vivo* (*6, 28, 29*), enabling us to study how nsp13 works against an opposing force of a DNA duplex and to determine what might occur in a head-collision with the RdRp. SPRNT thus provides a valuable platform for investigating nsp13’s role within the SARS-CoV-2 RTC.

Both translocation and unwinding with nsp13 appears to be strongly dependent upon the underlying DNA sequence. Several kinetic steps of the inchworm mechanism appear to be affected by the underlying DNA sequence. For example, during dsDNA unwinding, the advance of domain 1A is rate limiting and sequence-dependent (Fig. S16), while during ssDNA translocation with ATPγS, hydrolysis of ATPγS is rate-limiting and sequence-dependent in a different manner (Fig. S17). Similar sequence-dependent behavior has been found in all other helicases measured with SPRNT (Hel308, Rep, PcrA) (*14, 19*) and is evident in optical tweezers data with UvrD (*20*). However, Cryo-EM structures of nsp13 with an ATP analog and RNA in concert with RdRP (PDB ID: 7RDY) shows no base-specific interactions between the protein and RNA (*6*). Another hypothesis by which a helicase could have sequence-dependent translocation kinetics may be that differing steric effects of bases (rather than their binding affinities) in the NA binding channel allosterically influence the helicase’s conformational microstate thereby affecting various kinetic rates. Viral genomes are also highly structured such that particular sequence elements may play an important role in nsp13 regulation. SPRNT experiments with nsp13 on native SARS-CoV-2 sequences could shed light on such interactions.

SPRNT’s ability to determine how nsp13 works also sheds light on how it can be stopped from working. ATPγS is not a viable drug candidate for nsp13 because it is a general ATPase inhibitor, but we use ATPγS here to demonstrate the power of SPRNT in examining the mechanisms of helicase inhibition in exquisite detail. Our measurements allowed us to develop a model of ATPγS inhibition of nsp13 in which there are three mechanisms of action: 1) reduced processivity, 2) by preventing domain 1A and 2A from coming together after nucleotide binding, and 3) by slowing the hydrolysis of ATPγS relative to ATP. Similar studies with nsp13 drug candidates could be used to precisely determine the mechanism of inhibition and aid in development of more effective inhibitors.

## Supporting information

Supplemental tables, figures, and discussion

## Acknowledgments

S.K.M., J.M.C., C.A.T., A.M.P., S.J.A., J.C., M.C.F., J.R.H., H.C.K., H.D.B., J.H.G. and A.H.L. were supported by the National Institutes of Health, National Human Genome Research Institute (grant number R01HG005115). K.J.M. is supported by a National Cancer Institute K00 Fellowship (grant number K00CA223018). T.M.K. was supported by the National Institutes of Health, National Institute of General Medical Sciences for funding (grant number R35GM130234).

