## Supplemental tables, figures, and discussion for "Inhibition of the SARS-CoV-2 helicase at single-nucleotide resolution"

### Supplement for Inhibition of SARS-CoV-2 helicase at single-nucleotide resolution.

#### Methods:

DNA constructs: DNA oligonucleotides were synthesized at Stanford University Protein and Nucleic Acid Facility and purified at their facility using Poly-Pak cartridges. The oligo sequences are shown in Supplementary Table 1. For nsp13 unwinding assays, DNA template and complement were annealed at relative molar concentrations of either 1:1.2 or 1:10 respectively.

Proteins: The M2-NNN-MspA [1] protein was custom ordered from GenScript. M2-NNN-MspA used in nanopore experiments is identical to wild-type MspA (AC: CAB56052.1) except for the following mutations: D90N/D91N/D93N/D118R/E139K/D134R. Nsp13 from SARS-CoV-2 (AC: MT121215.1) was expressed and purified as reported previously [2].

Pore establishment: A single MspA nanopore was established in a bilayer as described previously [3]. Briefly, a lipid bilayer is formed with a mixture of 1,2-di-O-phytanyl-sn-glycero-3-phosphocholine (Avanti Polar Lipids, SKU: 999984C) and n-hexadecane across a 20 μm diameter aperture that separates two ~65 μL chambers (*cis and* trans) containing our operating buffers. A potential of 180 mV (unless otherwise specified for force titration experiments) is applied using an Axopatch 200B or Axopatch 1B amplifiers (Axon Instruments) and a National Instruments PCI-6251 DAQ.

Nanopore experiments:

All experiments were run at asymmetric salt, cis 20 mM KCl, with 20 mM HEPES at pH 7.5, 5 mM MgCl2 and 1 mM TCEP. The trans buffer 500 mM KCl, 20 mM HEPES at pH 7.5. Cis buffer also contained ATP, ADP, or ATPγS at variable concentrations. Nsp13 was preincubated with the DNA construct in the absence of ATP by incubating at RT for 30 min, at a final concentration of 4 µM and 0.4 µM respectively in 124 mM KCL, 20 mM Hepes pH 7.5, 5 mM MgCl2, and 1 mM TCEP. Unwinding and translocation experiments were initiated by adding Nsp13 and DNA to this cis well at final concentrations of 100nM and 10 nM respectively. Reagents were reperfused every 5 min to maintain constant concentrations of [ATP] and prevent bulk accumulation of [ADP] and [DNA]. All experiments were recorded at RT (23 $\pm$ 1 °C), with a bias of +180 mV (unless noted otherwise). ATP and TCEP was ordered from Sigma Aldrich and ATPγS were ordered from Sigma Aldrich or Tocris.

Data acquisition and analysis: Data were acquired with a sampling rate of 50 kHz and filtered at 10 kHz. Enzyme events were detected via thresholding: an event starts when ion current drops below 80% of the open pore current and ends after the ion current has returned above 94% of the open pore current for more than 8 datapoints. Candidate events are then evaluated by hand for processive ion-current leveling. Ion current steps within each event were found automatically using a point-by-point alignment of raw ion-current traces to a consensus of ion current levels (Fig. S5, and associated discussion). Event counts and statistics are summarized in Supplementary Table 2.

#### Table 1. DNA sequences used in this study:

Nsp13 experiments used template and complement sequences. All sequences are written 5′ – 3′.

| Strand Name | Strand Sequence |
|---|---|
| Template | PTTTTTTTTTTTTTTTTTTTTTTTTTTTTTTGAGTCCAAGTCCTGGACCTTTTGCCGAGCCGAGCCTTTTTTTTTTTTTTTTTTTTTTTTTTTTTTZ |
| Complement | GGCTCGGCTCGGCAAAAGGTCCAGGACTTGGA |

P = Phosphate, Z = Cholesterol tag.

#### Table 2. Summary of experimental statistics

A summary of the statistics of all the experiments performed in this study. All nsp13 experiments were conducted at a temperature of $23 \pm 1$ °C. The first column lists the experiment description.

| Experiment description | DNA substrate | [ATP] (μM) | [ATPγS] (μM) | [ADP] (μM) | Voltage (mV) | Number of events | Number of steps |
|---|---|---|---|---|---|---|---|
| **Translocation** | | | | | | | |
| Force dependence | ss | 3000 | -- | -- | 120 | 70 | 1254 |
| Force dependence | ss | 3000 | -- | -- | 180 | 2284 | 54785 |
| Force dependence | ss | 3000 | -- | -- | 220 | 86 | 1830 |
| ATP titration | ss | 1000 | -- | -- | 180 | 825 | 10992 |
| ATP titration | ss | 600 | -- | -- | 180 | 162 | 2251 |
| ATP titration | ss | 75 | -- | -- | 180 | 381 | 3576 |
| ATPγS titration | ss | 600 | 300 | -- | 180 | 258 | 2886 |
| ATPγS titration | ss | 0 | 600 | -- | 180 | 53 | 414 |
| ADP titration | ss | 600 | 0 | 600 | 180 | 445 | 10160 |
| ADP titration | ss | 75 | 0 | 600 | 180 | 84 | 929 |
| **Unwinding -- Template:complement 1:10** | | | | | | | |
| High ATP | ds | 1000 | -- | -- | 180 | 903 | 9427 |
| **Unwinding -- Template:complement 1:1.2** | | | | | | | |
| ATPγS titration | ds | 600 | 0 | -- | 180 | 428 | 8601 |
| ATPγS titration | ds | 600 | 150 | -- | 180 | 54 | 293 |
| ATPγS titration | ds | 600 | 300 | -- | 180 | 263 | 1374 |
| ATPγS titration | ds | 600 | 600 | -- | 180 | 260 | 1414 |
| ATPγS titration | ds | 0 | 600 | -- | 180 | 154 | 701 |
| ATPγS titration, force | ds | 600 | 300 | -- | 120 | 45 | 511 |
| ATPγS titration, force | ds | 600 | 300 | -- | 150 | 61 | 586 |
| ATPγS titration, force | ds | 600 | 300 | -- | 220 | 137 | 1280 |

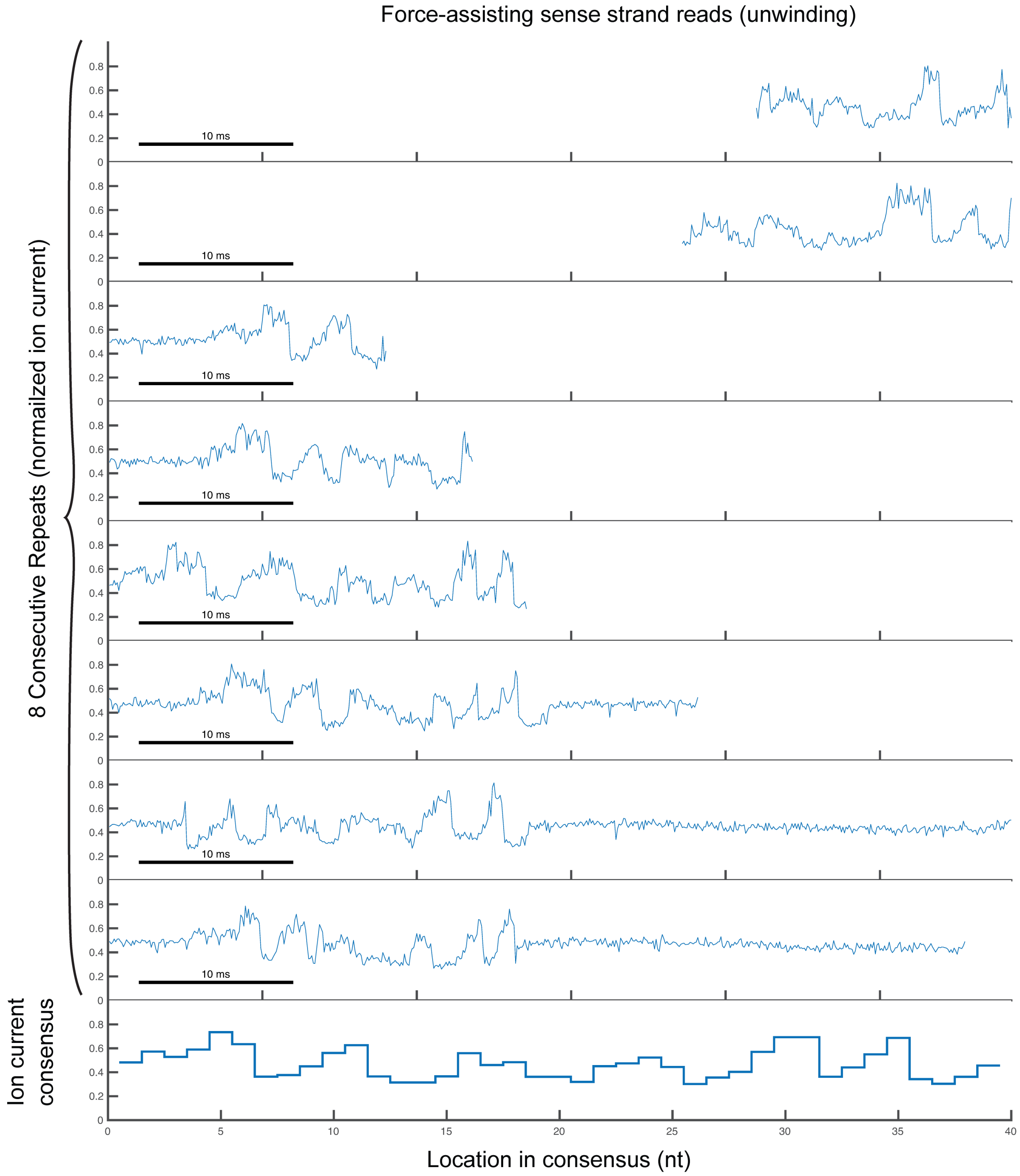

#### Figure S1: Example data traces for nsp13 translocation on ssDNA

Eight example data traces recorded at 180mV (36 pN) displayed at 10kHz sampling rate. The bottom graph shows the empirically determined ion-current consensus for the template strand. Levels within the ion current trace are automatically detected and aligned to the ion current consensus using a point-by-point alignment algorithm previously described.

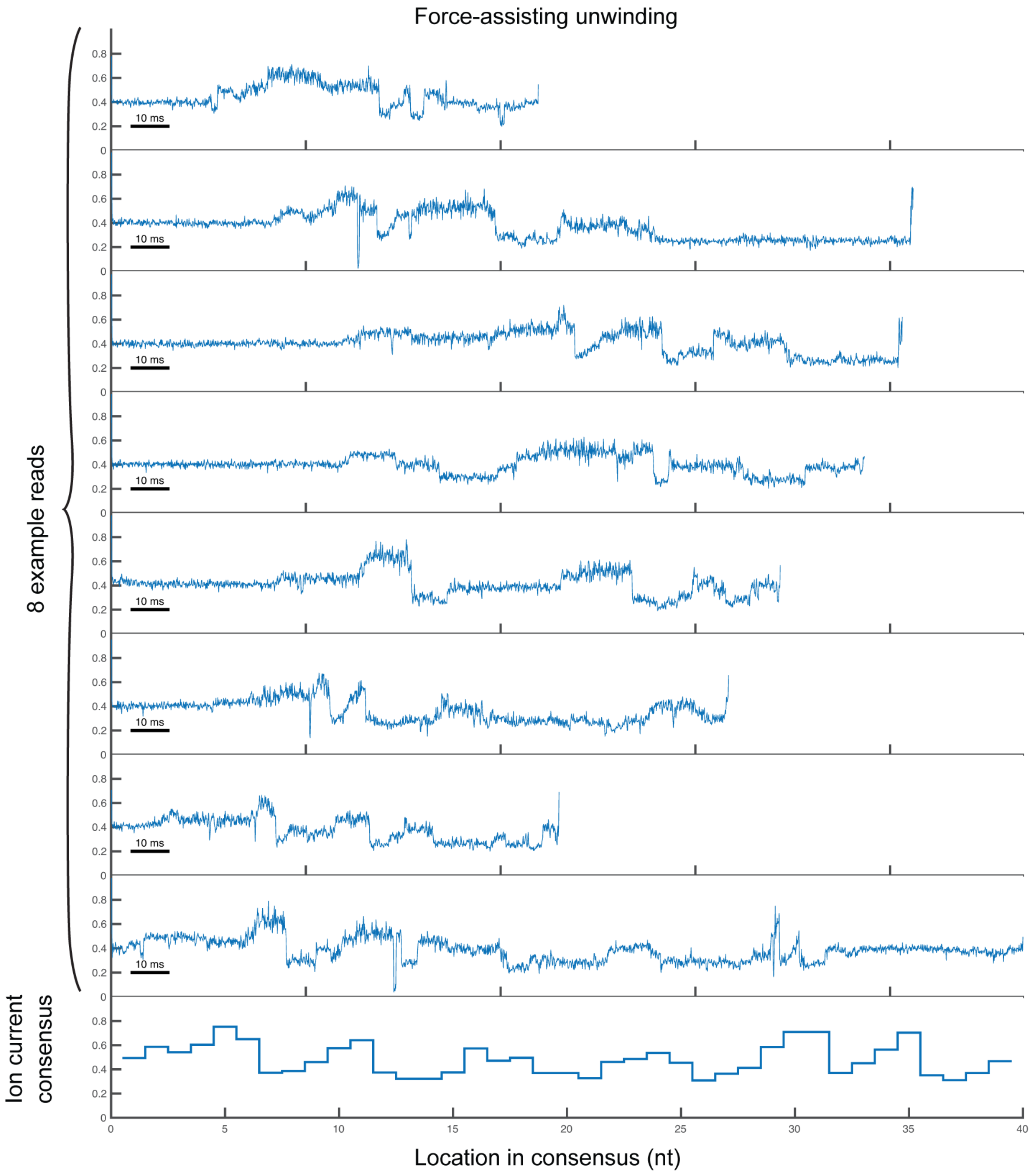

#### Figure S2: Example data traces for nsp13 unwinding on dsDNA

Eight example data traces recorded at 180mV (36 pN) displayed at 10kHz sampling rate. The bottom graph shows the empirically determined ion-current consensus for the template strand. Levels within the ion current trace are automatically detected and aligned to the ion current consensus using a point-by-point alignment algorithm previously described.

#### Figure S3: Schematic of nsp13 resting on-top of MspA nanopore

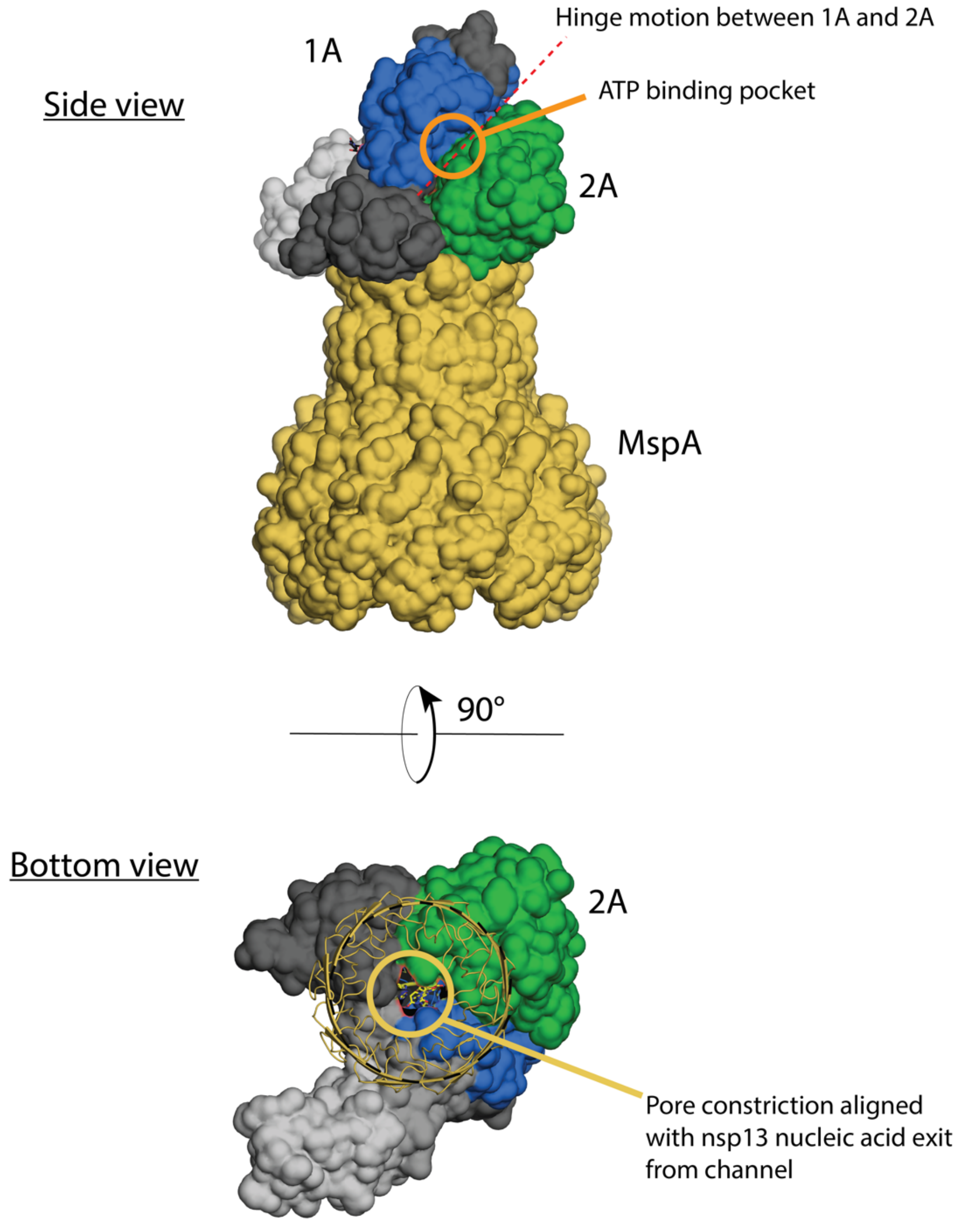

Schematic of nsp13 feeding DNA into the nanopore During SPRNT, enzyme-NA complexes are electrophoretically drawn into the nanopore until the enzyme comes to rest on the rim of the pore. In force-assisting measurements, enzyme is lowering DNA in the same direction as the electrostatic force on the DNA. Using cryo-EM structure of nsp13 bound with RNA (PDB ID: 7RDY), we orient the enzyme on MspA (PDB ID: 1UUN) such that 5′-end of RNA is fed into the pore. This would result in domain 2A directly in contact with MspA. The applied force of ~36 pN acts on domain 2A in the same direction as enzyme motion from 5′-end towards domain 1A at the 3′-end.

#### Figure S4: Speed distributions for NSP13 translocation and unwinding

Probability distributions of enzyme speed for ssDNA translocation at 75 μM, 600 μM, and 1000 μM ATP and dsDNA unwinding at 1000 μM ATP (a – d respectively). The histograms are constructed by taking all measurements of speed of enzymes (1/ mean duration per step per event; reciprocal of the mean dwell-time per position of each enzyme event). The number of events recorded in each condition is displayed above the distribution.

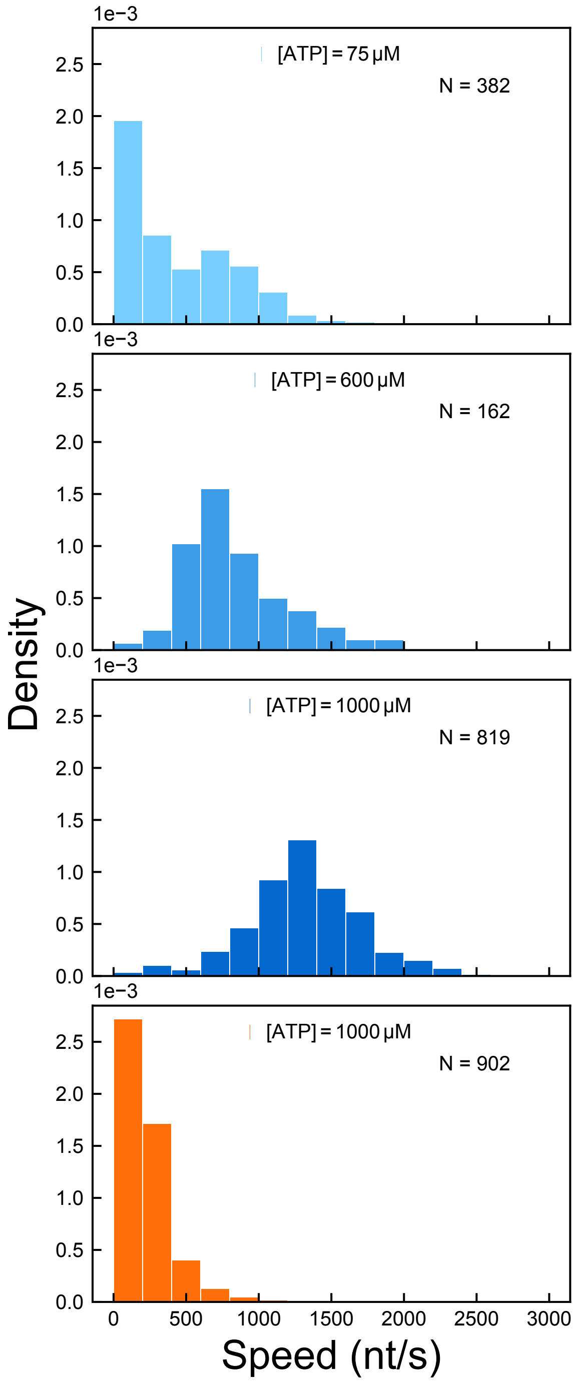

### Figure S5: Sequence-aware step detection

A

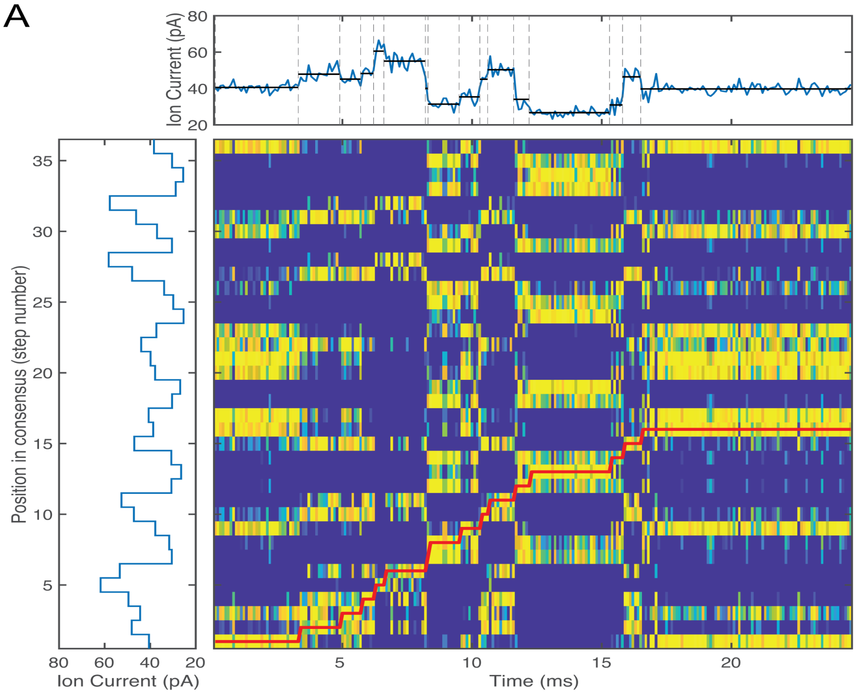

B

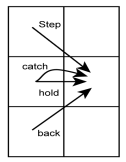

C

$$A_{n,m} = \max \begin{cases} A_{n+1,m-1} + \log(p_{step}) + S_{n,m} \\ A_{n+1,m} + \log(p_{hold}) + S_{n,m} \\ A_{n+1,m+1} + \log(p_{back}) + S_{n,m} \\ A_{n+1,m} + \log(p_{catch}) \end{cases}$$

D

| | |
|---|---|
| $\log(p_{step})$ | -5.0 |
| $\log(p_{hold})$ | -0.007 |
| $\log(p_{back})$ | -7.4 |
| $\log(p_{catch})$ | -16.8 |

A)Top: an example raw ion-current trace for nsp13 translocating on ssDNA. Left: Ion-current consensus for the template strand. For alignment, gaps in the consensus that occur due to missing ion-current levels are omitted. Aligned position is later corrected by inserting gaps where they occur. The ion-current consensus was built via comparison of several nsp13 dsDNA unwinding reads to ion-currents predicted by the previously measured ion-current-to-sequence map. Middle: A similarity matrix comparing each datapoint in the raw data to the consensus shown at left. Warmer colors indicate close agreement with the consensus current values while cool colors indicate significant differences. Score values are calculated as $S_{n,m} = -0.5*\ln(2\pi*V)+(X_m-R_n)/V$. Where V is the average current noise squared, $X_m$ is the mth datapoint and Rn is the nth consensus current level. Alignment of the currents in X to the consensus values of R, (shown in red), uses the similarity matrix along with the allowed step types shown in B). Steps within the alignment can then be used to determine the most likely location of an ion-current step (top: gray dashed lines) and to locate ion current levels within the raw ion current trace (top: black lines) B). Alignment proceeds via a modified version of the dynamic-programming alignment scheme described in[4]. C) The first column in A is initialized with the first column of the similarity score. From there each element of the alignment matrix, A, is filled out individually by choosing the best score for the given step type. The “catch” option deals with occasional spurious noise spikes allowing the alignment to ignore brief deviations in ion-current that are not described well by the consensus. However, because the catch penalty is large, it is rarely invoked by the algorithm. D) Table of values used for the various step penalties. These values are log probabilities and can be tuned to adjust for differing step durations, however alignment is robust to detuning of step penalties. For example, here, $p_{hold} = e^5 * p_{step} \sim 150 * p_{step}$, indicating an expected ~150 datapoints (15 ms) per step. However, the alignment still performs well on steps far shorter than 15 ms in duration. This can be attributed to the relatively high signal-to-noise ratio of the data. There is really only one path through the similarity matrix which does not step through a region of cool colors. Alignment algorithms such as this provide a wholistic alignment of the data. The alignment path passing through one region of the similarity matrix requires that it can continue on through the subsequent regions. For example a datapoint at 14 ms into the event matches reasonably well with consensus levels 13, 19, 24, 34, but is aligned to consensus level 13 not only because it matches well, but also because all the other datapoints before and after it match the other surrounding levels. Similar alignments are regularly used for biological sequence alignment such as alignment of DNA sequences and is guaranteed to find the optimal alignment of X to R for the given set of rules[5].

### Figure S6: Alignment of ssDNA translocation traces

Example alignments of 8 representative ssDNA translocation reads at [ATP] = 1000μM. The representation is the same as is shown in Figure 1. At the top of each graph, there is a representation of the ion current consensus to which the raw data is aligned. In the middle is the raw ion-current data for the read and at the bottom is the aligned sequence position as a function of time. Ion current is displayed at 10kHz. Red dashed lines at positions 5, 10, 15, and 30 serve as guides to the eye to aid in visual verification of the fidelity of the alignments. Step transitions and dwell-times can then be readily extracted from the alignment information.

ssDNA [ATP] = 1000 μM

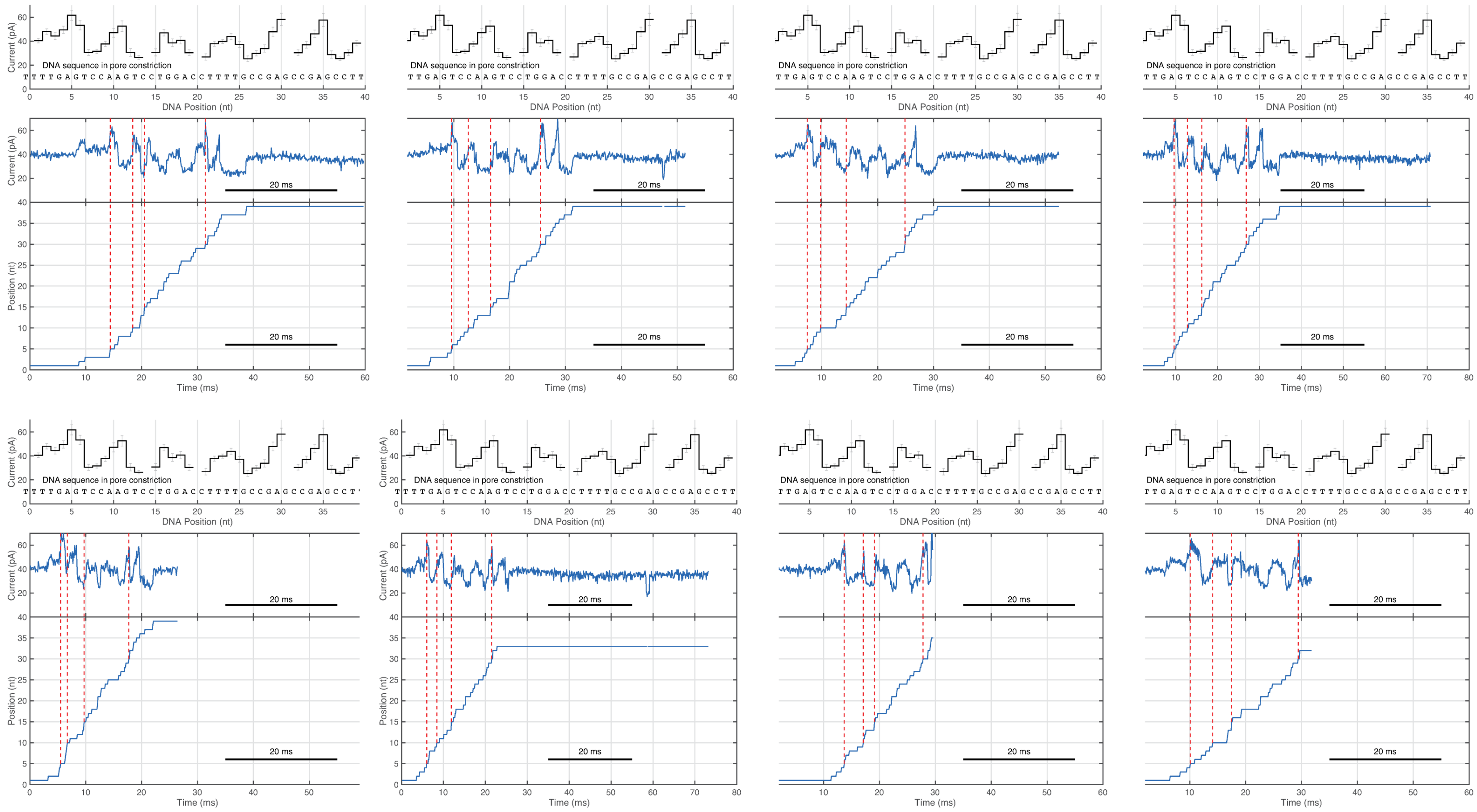

#### Figure S7: Alignment of ssDNA translocation traces

Example alignments of 8 representative dsDNA unwinding reads at [ATP] = 1000μM. The representation is the same as is shown in Figure 1. At the top of each graph, there is a representation of the ion current consensus to which the raw data is aligned. In the middle is the raw ion-current data for the read and at the bottom is the aligned sequence position as a function of time. Ion- current is displayed at 10kHz. Red dashed lines at positions 5, 10, 15, and 30 serve as guides to the eye to aid in visual verification of the fidelity of the alignments. Step transitions and dwell-times can then be readily extracted from the alignment information.

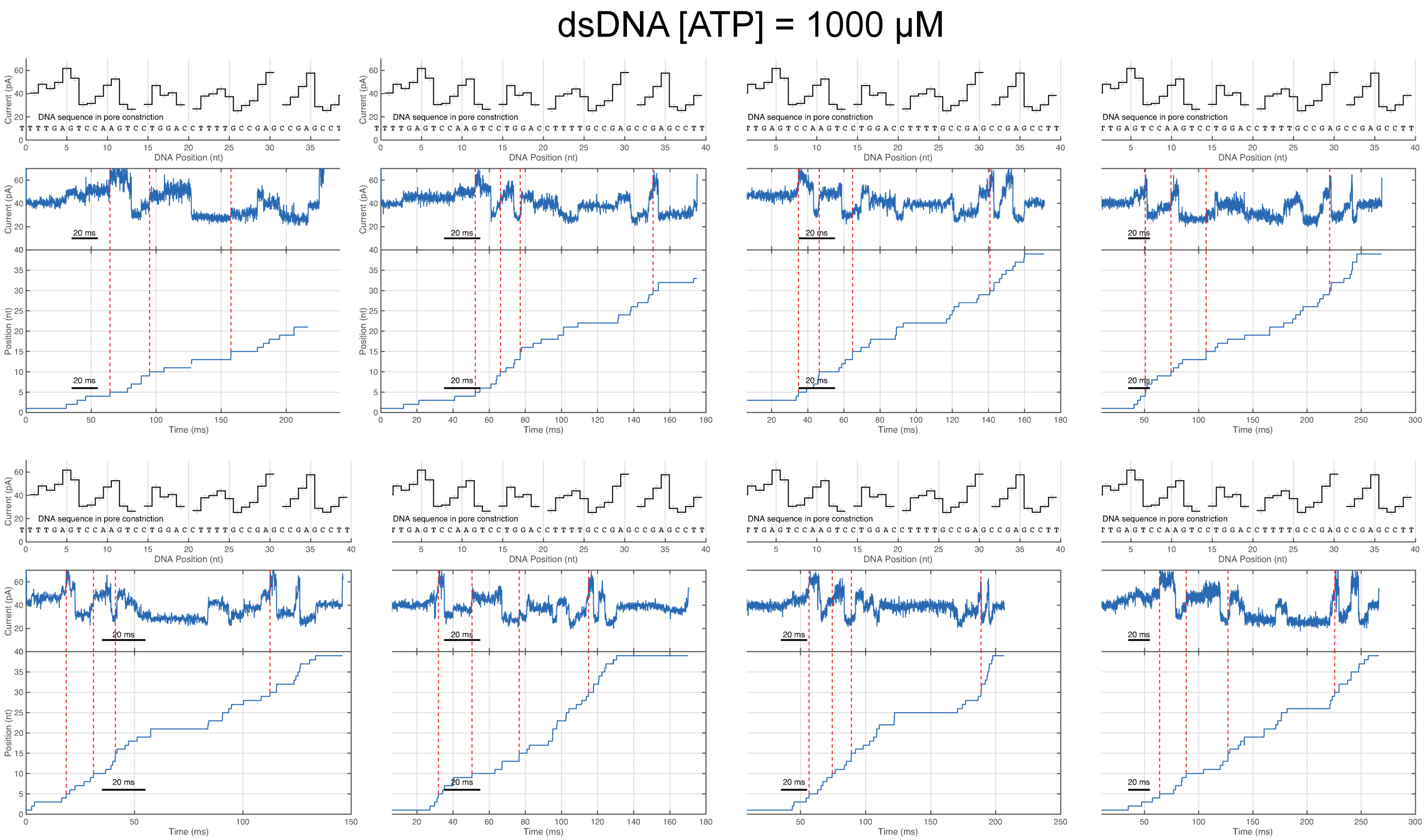

#### Figure S8: Dwell-time distributions for each step at 1000 µM ATP and 75 µM ATP in ssDNA translocation

Probability distributions for the durations of individual levels from Figure 3C (left). (A) at saturating [ATP] and (B) low (rate-limiting). The y-axis is logarithmic. The histograms were constructed by taking all measurements of the dwell-times of a given level at one [ATP] condition. The red line is the maximum likelihood exponential fit to the data and the orange line indicates the mean of the distribution. Most data sets are well described by a single exponential. The mean dwell-time ± S.E.M. (1 sigma confidence intervals) are displayed with each distribution. The mean dwell-time varies from level to level in a statistically significant way, possibly due to sequence dependence.

A

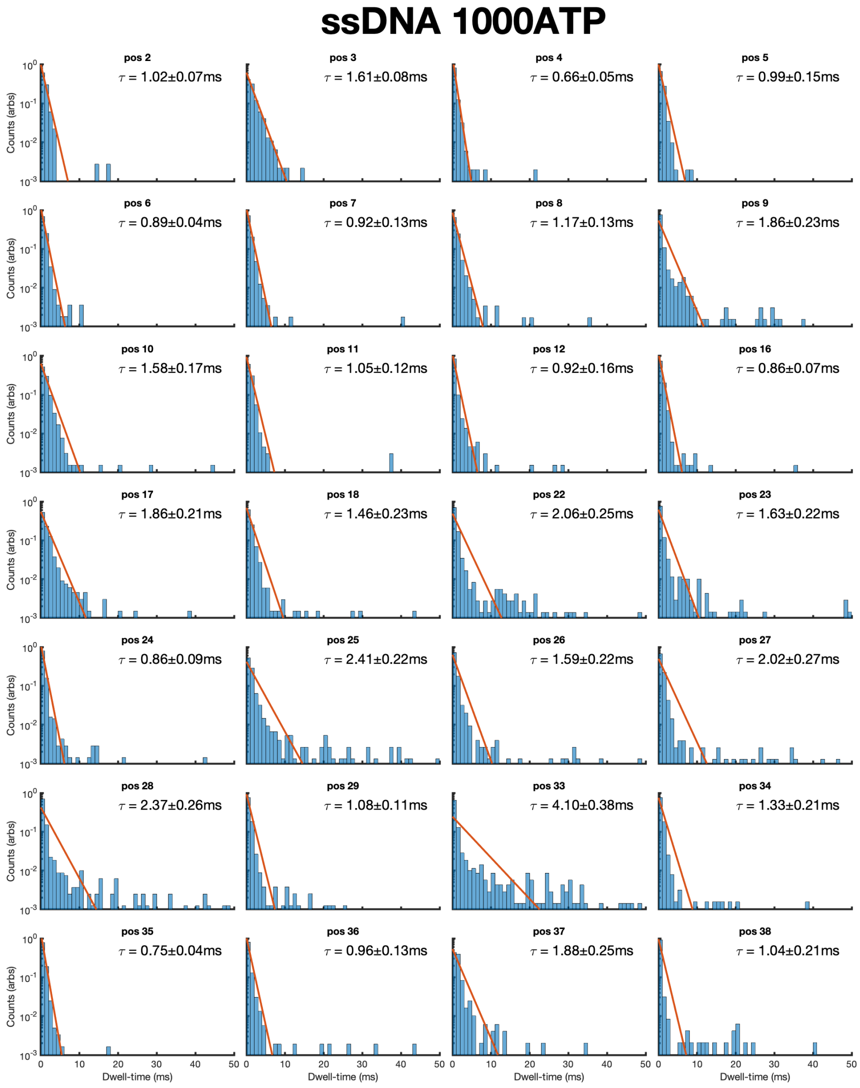

B

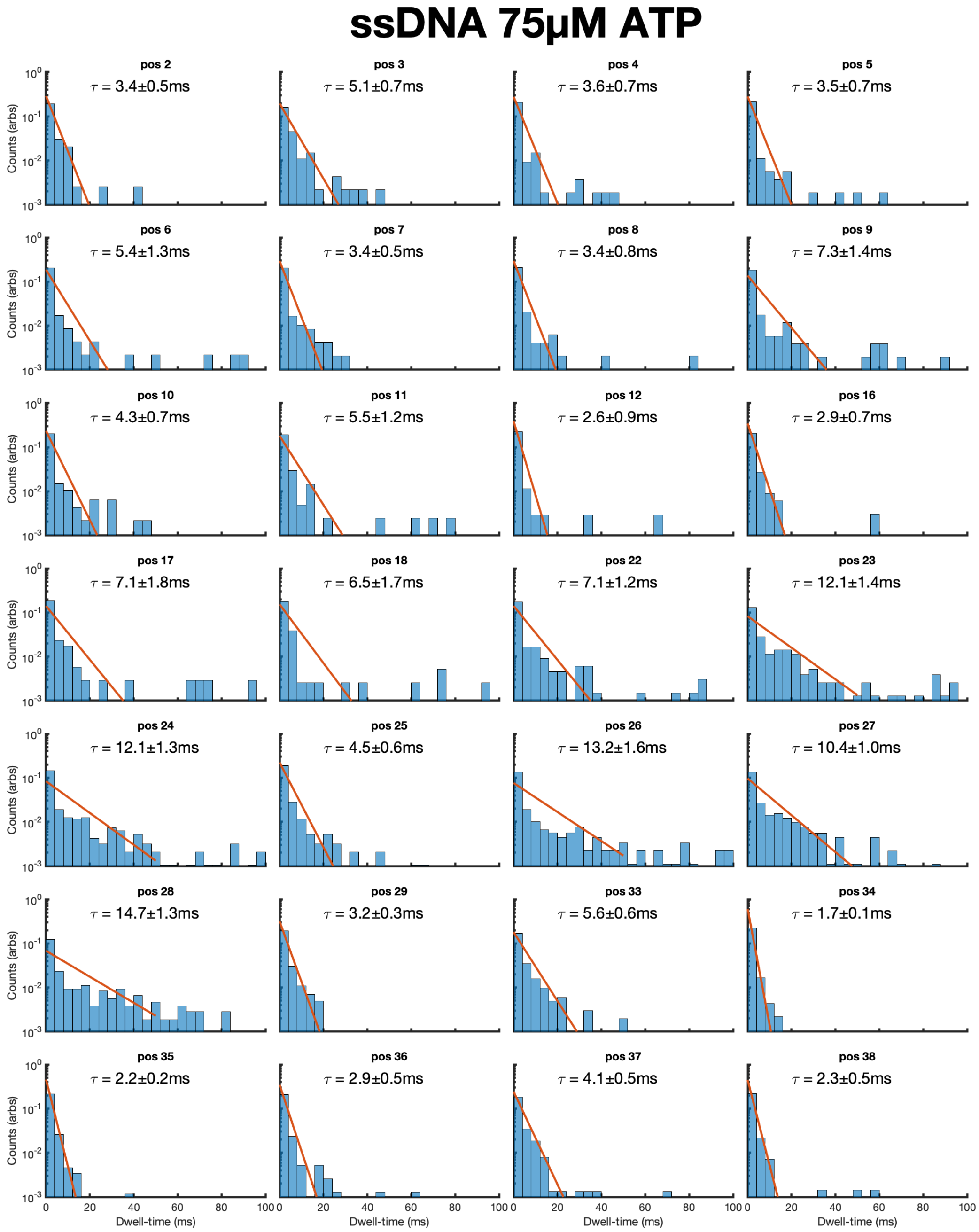
ssDNA 75µM ATP
pos 2
τ = 3.4±0.5ms
pos 3
τ = 5.1±0.7ms
pos 4
τ = 3.6±0.7ms
pos 5
τ = 3.5±0.7ms
pos 6
τ = 5.4±1.3ms
pos 7
τ = 3.4±0.5ms
pos 8
τ = 3.4±0.8ms
pos 9
τ = 7.3±1.4ms
pos 10
τ = 4.3±0.7ms
pos 11
τ = 5.5±1.2ms
pos 12
τ = 2.6±0.9ms
pos 16
τ = 2.9±0.7ms
pos 17
τ = 7.1±1.8ms
pos 18
τ = 6.5±1.7ms
pos 22
τ = 7.1±1.2ms
pos 23
τ = 12.1±1.4ms
pos 24
τ = 12.1±1.3ms
pos 25
τ = 4.5±0.6ms
pos 26
τ = 13.2±1.6ms
pos 27
τ = 10.4±1.0ms
pos 28
τ = 14.7±1.3ms
pos 29
τ = 3.2±0.3ms
pos 33
τ = 5.6±0.6ms
pos 34
τ = 1.7±0.1ms
pos 35
τ = 2.2±0.2ms
pos 36
τ = 2.9±0.5ms
pos 37
τ = 4.1±0.5ms
pos 38
τ = 2.3±0.5ms
Counts (arbs)
Dwell-time (ms)

#### Figure S9: ATP titration of nsp13 helicase by DNA position

Enzyme rate as a function of [ATP] at several DNA positions. Rate is calculated as the reciprocal of mean dwell-time at the DNA position. The mean dwell-time at each [ATP] was calculated by taking the average of all dwell-times from t=0 s to t=100 s. The black line shows the best fit of the data to the Michaelis-Menten equation. The red and blue line correspond to maximum velocity of reaction ($V_{max}$) and Michaelis constant ($K_m$) for ATP.

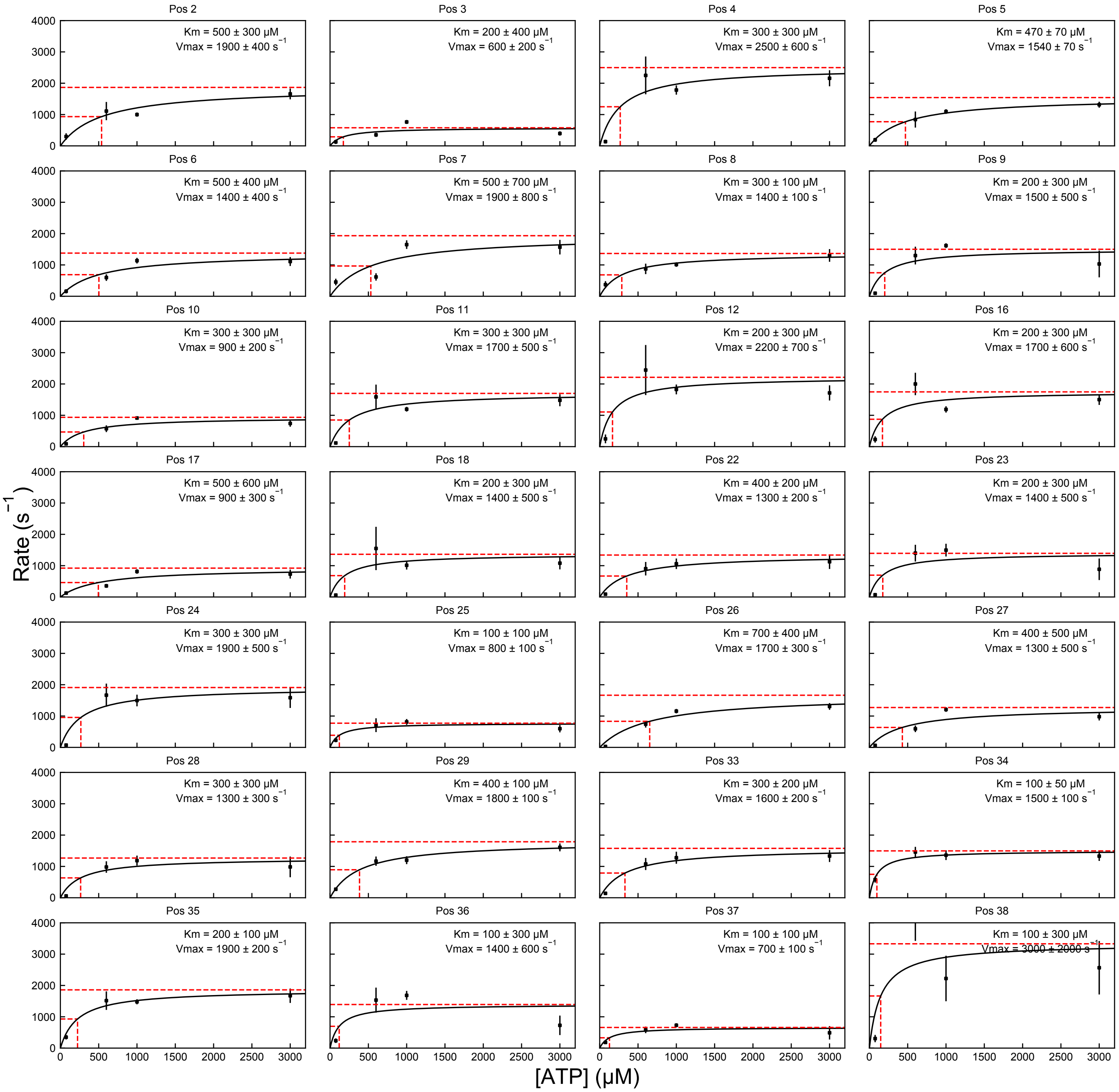

#### Figure S10: Read start and end location histogram (ssDNA and dsDNA)

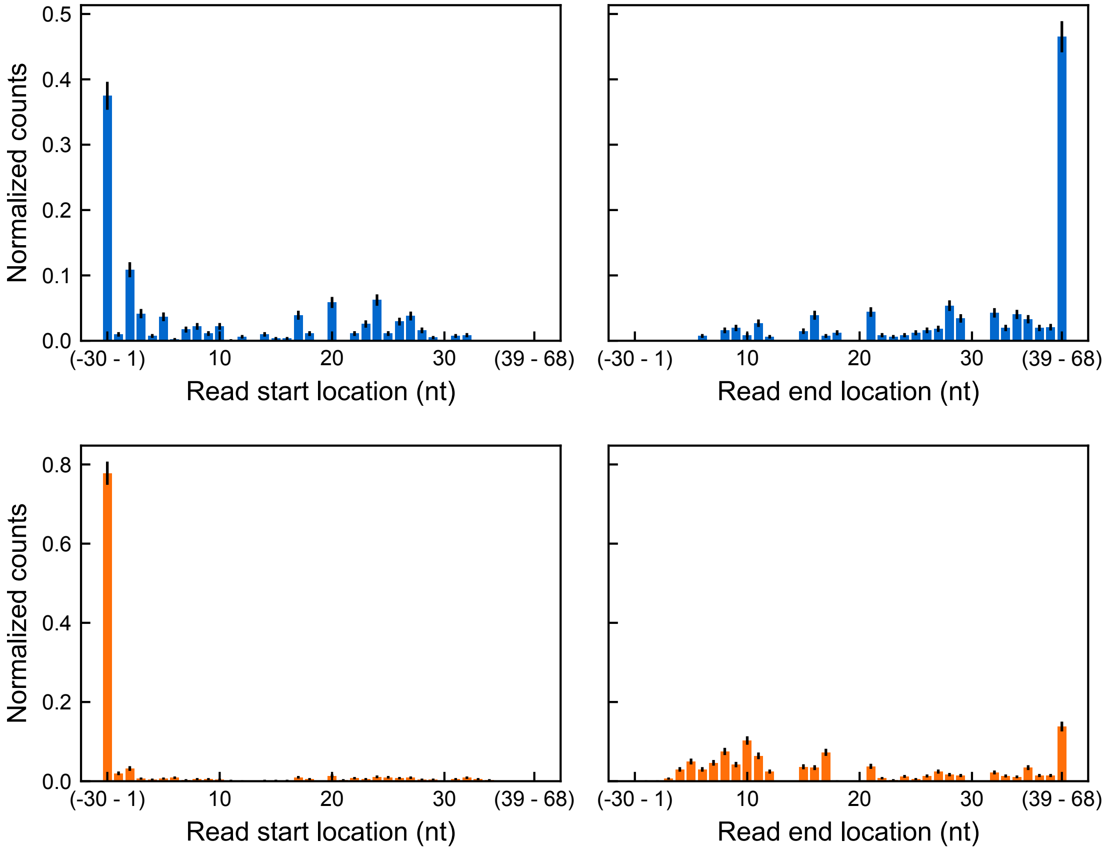

**Figure S11: Effect of force on nsp13 translocation kinetics**

The effect of applied force on nsp13 translocation on ssDNA was studied by varying the voltage at constant saturating [ATP]=3 mM (A) Probability distributions of dwell-time (ms) measured at 3 different voltages 220 mV, 180 mV, 120 mV that corresponds to ~44 pN, ~36 pN, ~24 pN force respectively. N_pores is number of independent trials and N_counts is the total number of positions measured in each condition. (B) Mean dwell-time (ms) (mean ± s.e.m) as a function of force (pN).

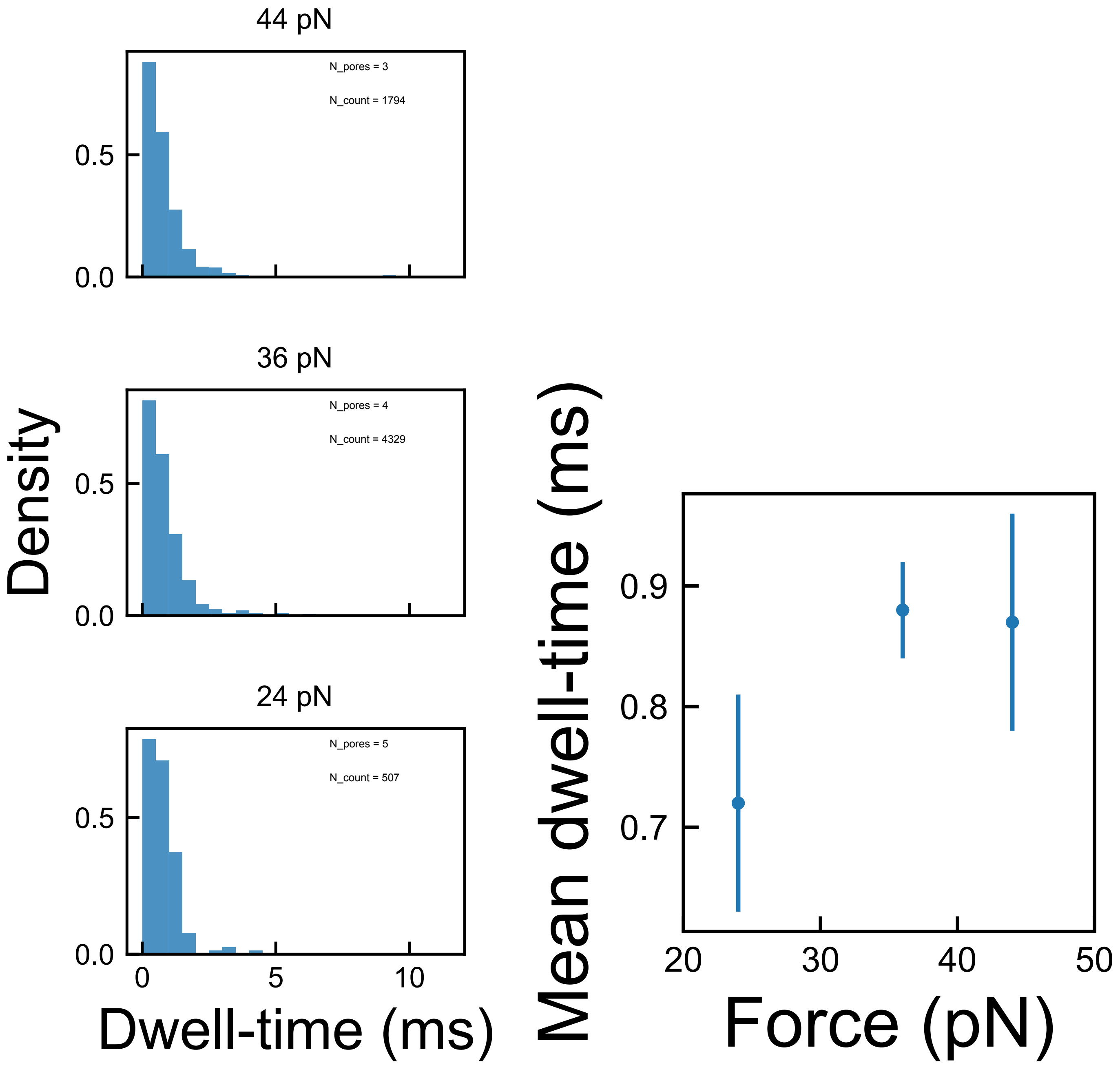

#### Figure S12: Effect of force on nsp13 translocation kinetics by nucleotide position

The data used in Fig. S11 broken down by position

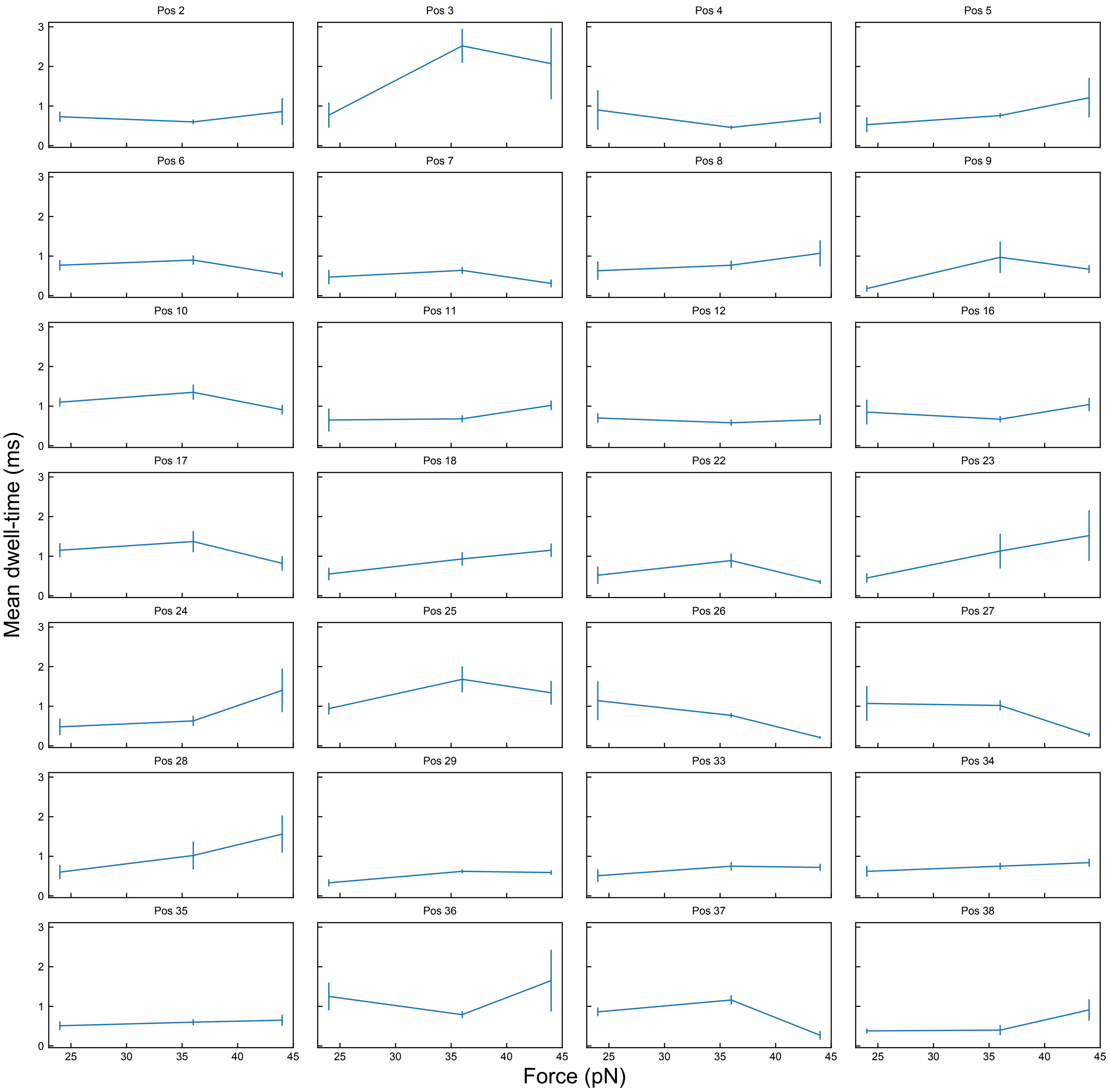

### Figure S13: Effect of ADP on nsp13 ssDNA translocation

S13A: Dwell-time histograms for ssDNA translocation of nsp13 helicase at each position along the DNA strand with 600µM ATP. Dwell-times are slightly longer than those observed at 1000µM (Fig. S8A). This graph is a point of comparison for data with 600µM ATP and 600µM ADP (Fig. S13B).

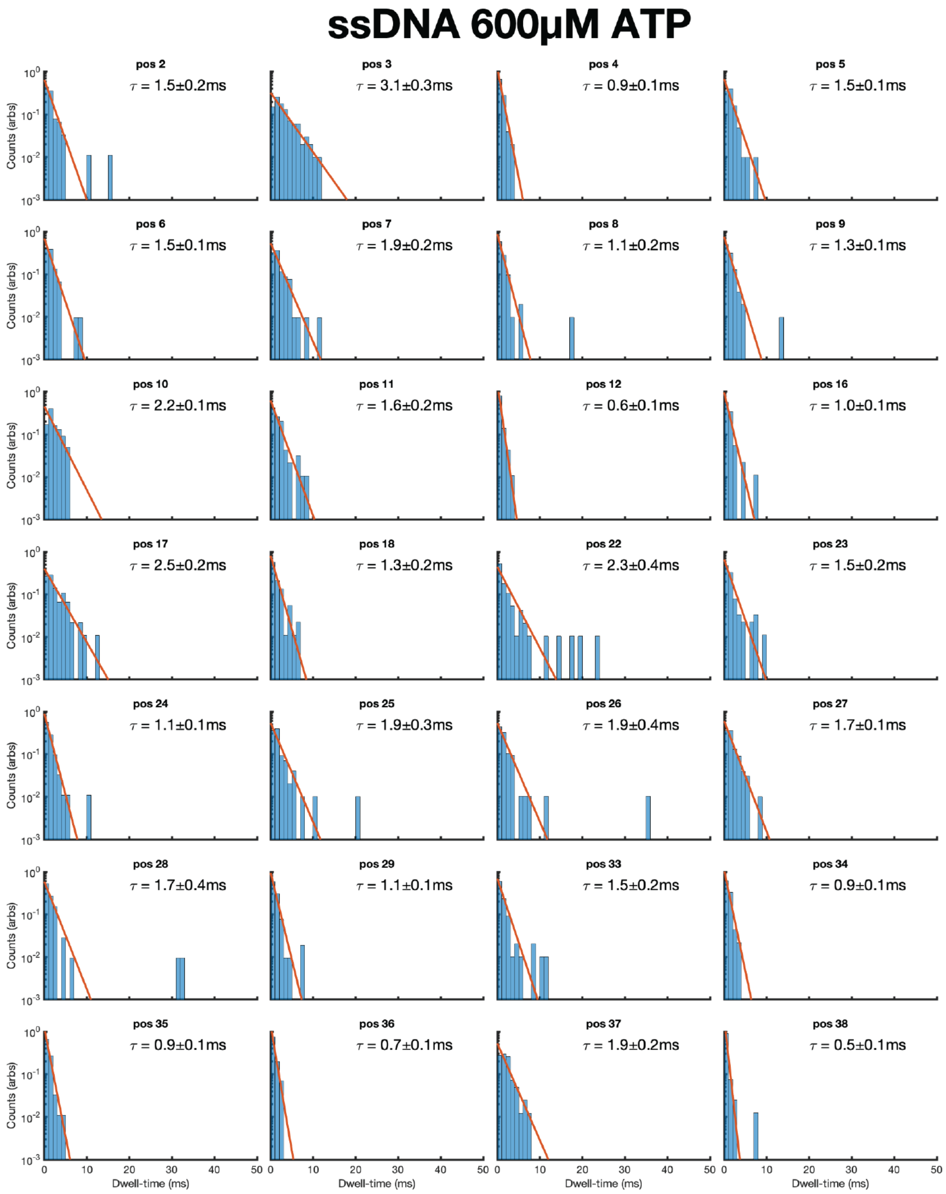

S13B: Dwell-time histograms for ssDNA translocation of nsp13 helicase at each position along the DNA strand with 600μM ATP and 600μM ADP. Dwell-times are ~2x longer than those observed at with 600μM ATP and are consistent with the competitive inhibition observed with other monomeric helicases using SPRNT[6,7].

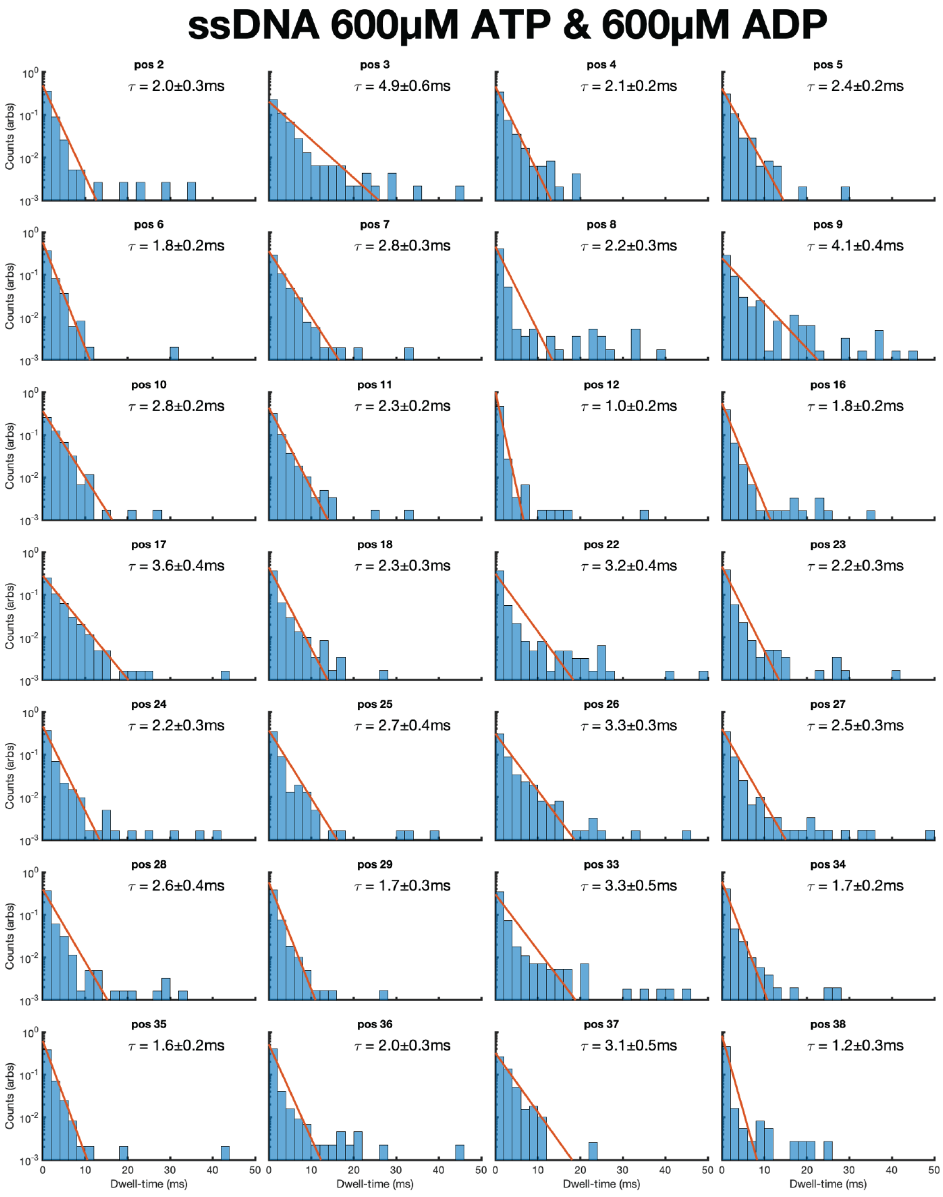

#### Figure S14: Effect of complement strand concentration on unwinding speed

(A) Probability distributions of enzyme speed in the presence of ssDNA (top), dsDNA with template and complement strands in 1:1.2 ratio (middle) and dsDNA with template and complement strands in 1:10 ratio (bottom). (B) Probability distributions of read length for corresponding nsp13 events and conditions in (A). (C) Read coverage graph showing individual nsp13 reads plotted with their start and end location (x-axis), as a percentile of their start location (y-axis). (D) Distributions of average dwell time per event (averaged across all positions) for events where the read started before position 5 (left) and after position 5 (right).

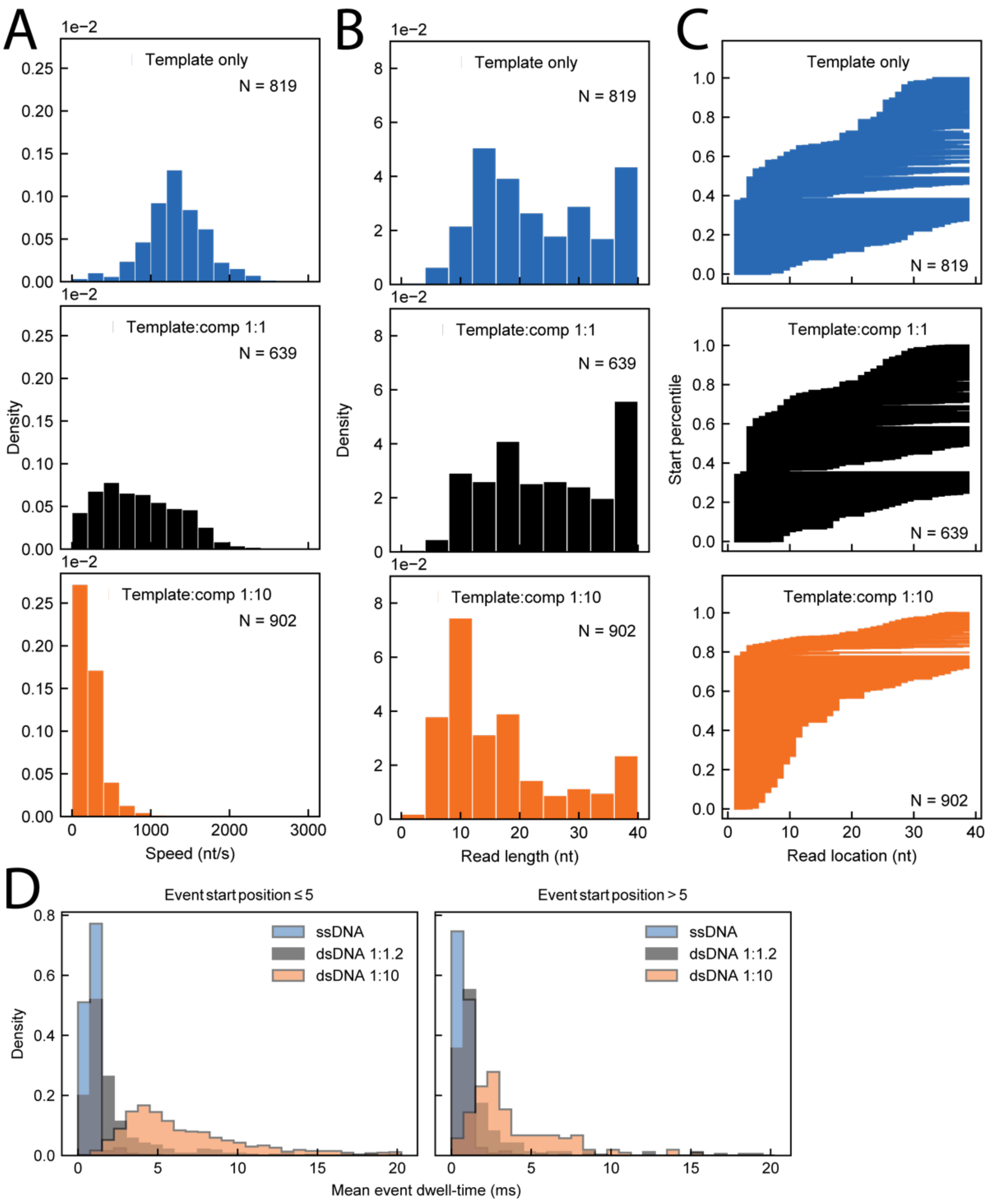

**Figure S15: Comparing event dwell-time distributions as a function of start location, [ATP] and substrate (dsDNA 1:1.2, dsDNA 1:10, ssDNA)**

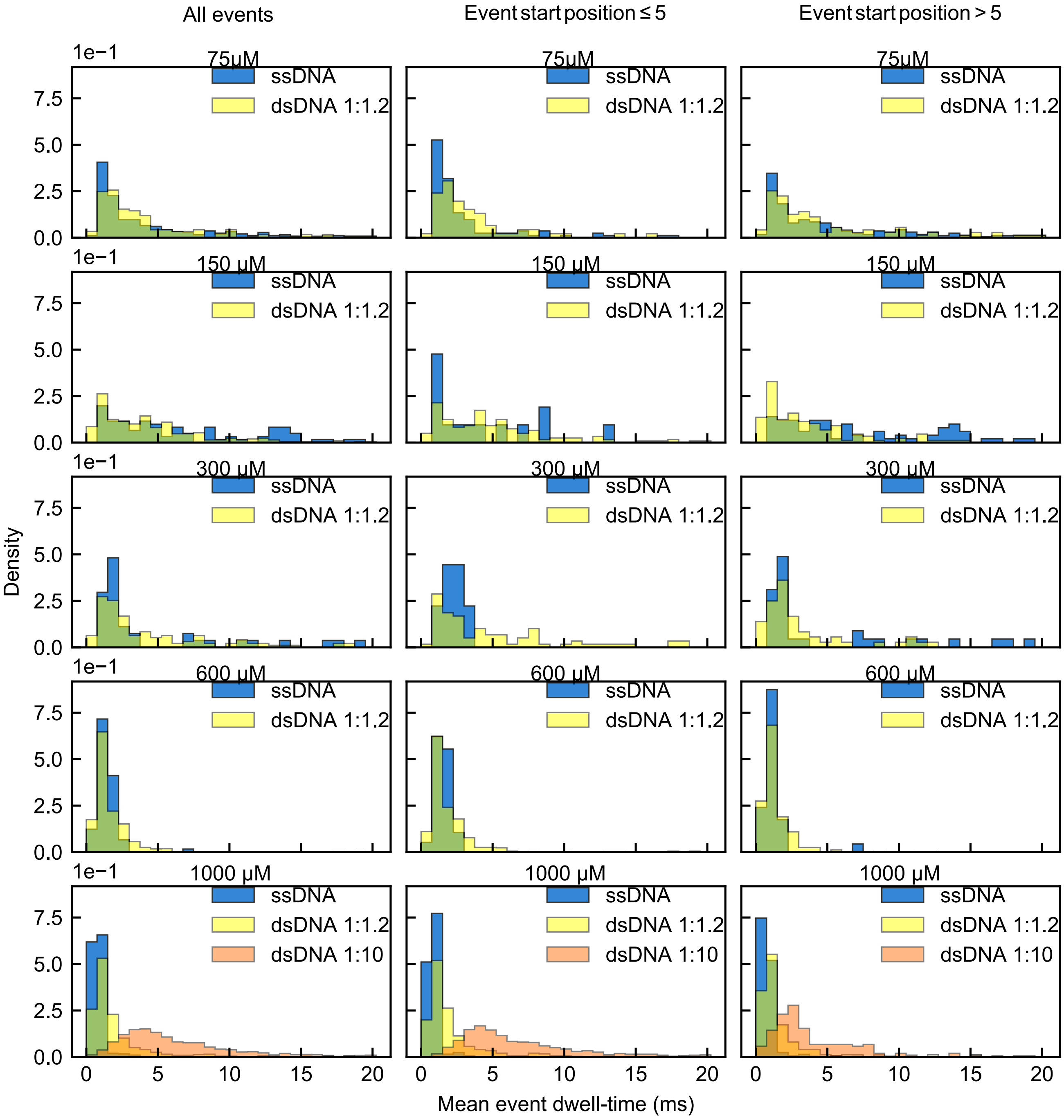

#### Figure S16: Dwell-time distributions for each step at 1000 µM ATP during dsDNA unwinding

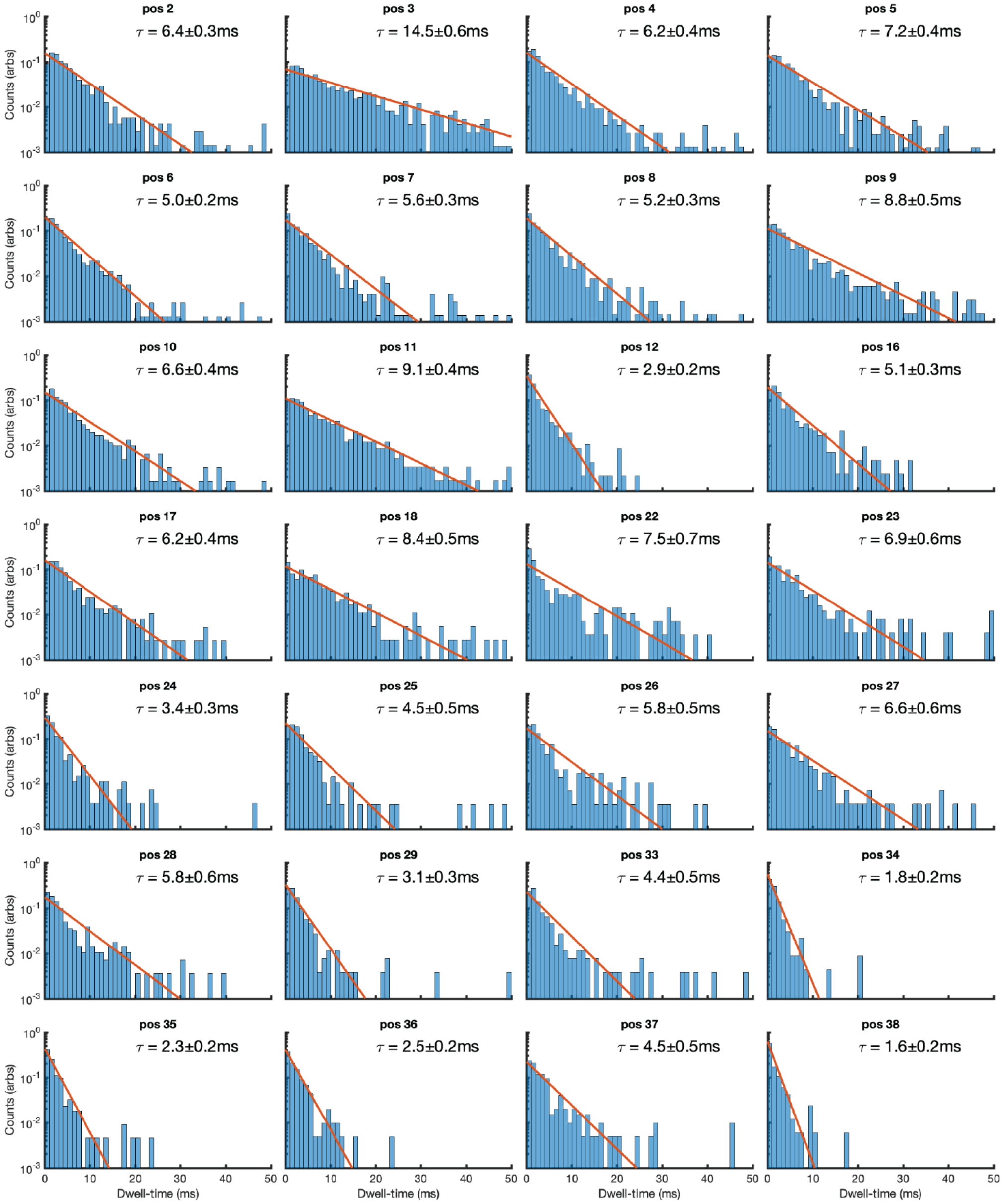

Probability distributions for the durations of individual levels in Figure 3C (right, dsDNA unwinding) at saturating [ATP]. The y-axis is logarithmic. The histograms were constructed by taking all measurements of the dwell-times of a given level at one 1000 µM ATP condition. The red line is the best fit exponential to the data. Most positions are well described by a single exponential. The mean dwell-time ± S.E.M. are displayed with each distribution. The dwell-time varies from level to level in a statistically significant way, possibly due to interactions between nsp13 and the underlying DNA sequence.

#### Figure S17: Dwell-time distributions for each step with ssDNA with 600μM ATPγS-only

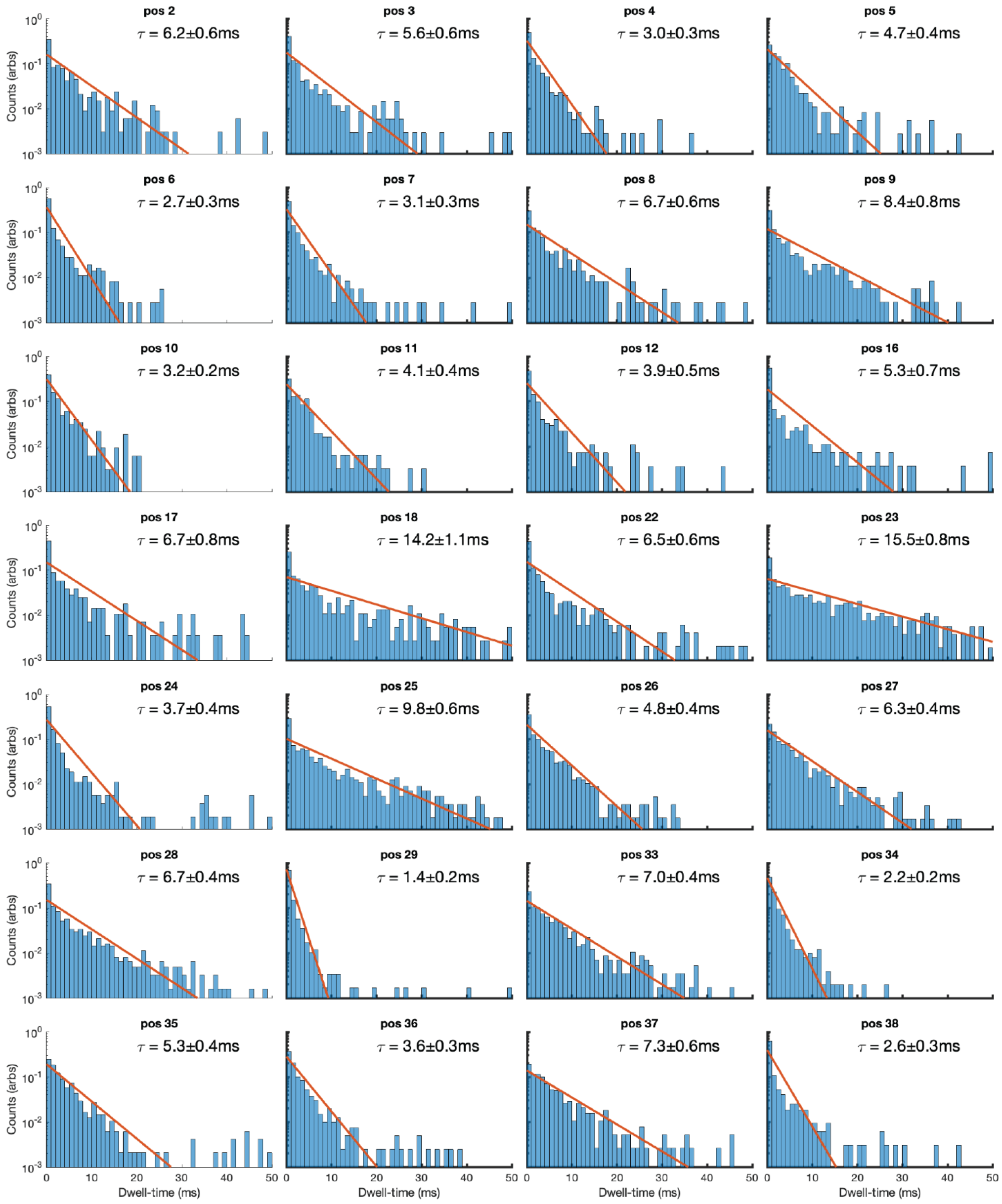

Probability distributions for the durations of individual levels with 600μM ATPγS-only. The y-axis is logarithmic. The histograms were constructed by taking all measurements of the dwell-times of a given level at one 1000 μM ATP condition. The red line is the best fit exponential to the data. Most positions are well described by a single exponential. The mean dwell-time ± S.E.M. are displayed with each distribution. The dwell-time varies from level to level in a statistically significant way, possibly due to interactions between nsp13 and the underlying DNA sequence. ATPγS clearly slows nsp13 down in comparison to ssDNA with saturating [ATP] (Fig. S8A). Intriguingly, the apparent sequence-dependence is different for ATPγS as compared to dsDNA. In other words, steps that are slowest for unwinding are not necessarily the slowest for translocation using ATPγS. In most locations, duplex unwinding is slower than ATPγS-hydrolysis though they are a similar order-of-magnitude.

### Figure S18: Alternative models for ATPyS inhibition of nsp13 unwinding

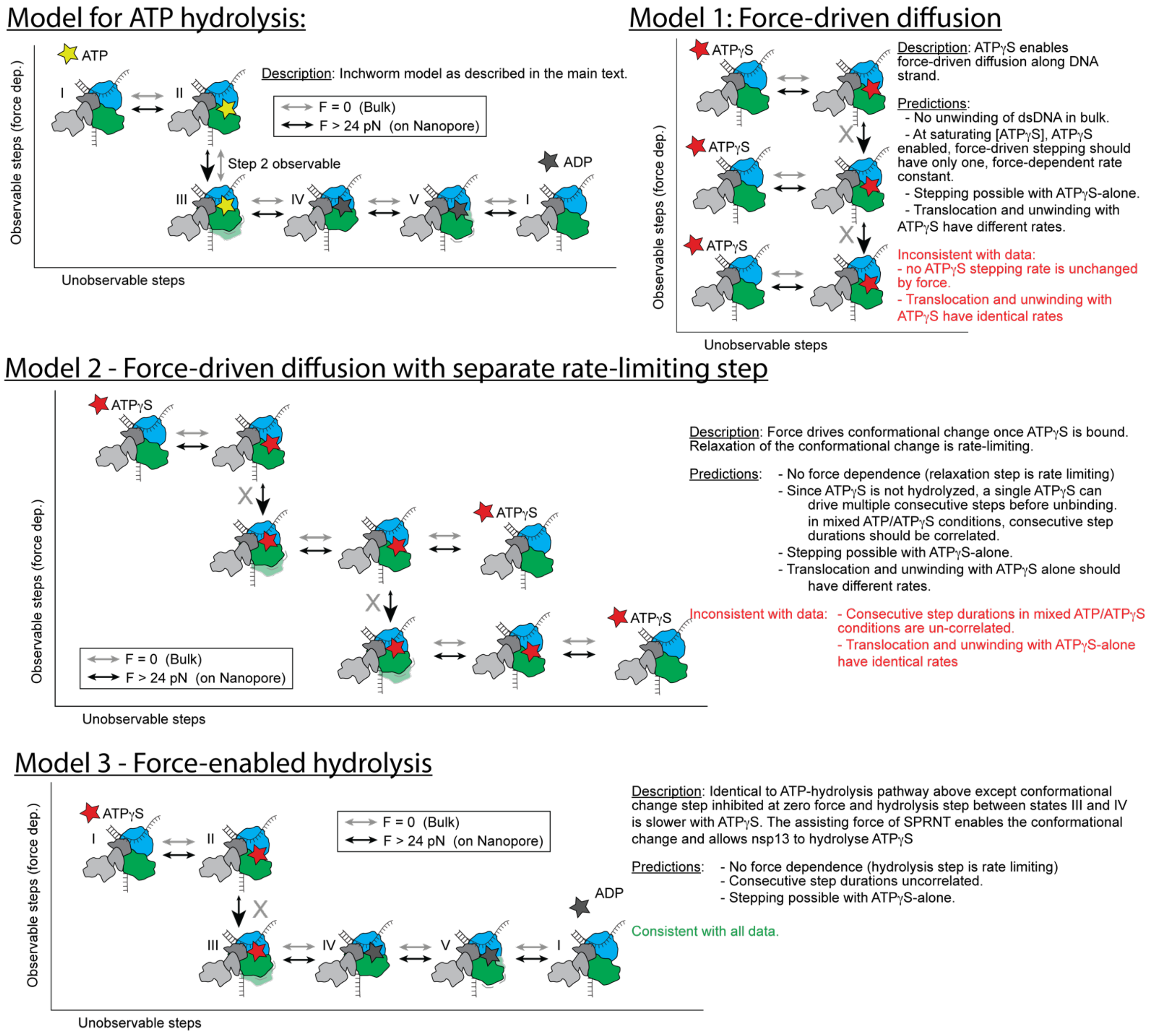

#### Figure S19: Lack of autocorrelation in ATPγS data dwell times

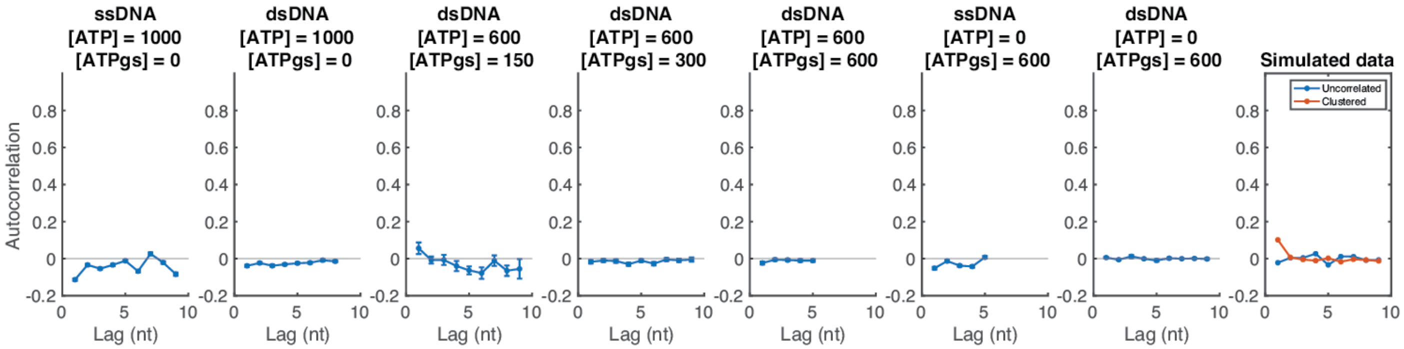

In an alternative model for inhibition in which ATPγS enables translocation but is not hydrolyzed (e.g. Models 1 or 2 in Fig. S18), adjacent step durations could be correlated in mixed ATP and ATPγS conditions. If ATP□S is not hydrolyzed, once an ATPγS is bound, it can conceivably be used for multiple consecutive steps. The above graphs show the autocorrelation as a function of various lags for the indicated experimental conditions, displayed concentrations are in µM. Autocorrelations of the log step durations for each single-molecule trace were calculated and then averaged over all molecules. If ATPγS-inhibited steps appeared in clumps in the data as is predicted by Models 1 and 2, we would expect to see an appreciable increase in the autocorrelation of dwell-times for conditions with mixed ATP and ATPγS in which ATP-driven and ATPγS-driven steps occur (e.g. simulated data in red in the rightmost plot). Instead, the autocorrelation plots for all conditions more closely resemble uncorrelated adjacent dwell-times (e.g. simulated data in blue in the rightmost plot). This effectively rules out models in which ATPγS enables translocation and unwinding but is not hydrolyzed.

#### Figure S20: Dwell-time distributions for each step with dsDNA with 600μM ATPγS-only

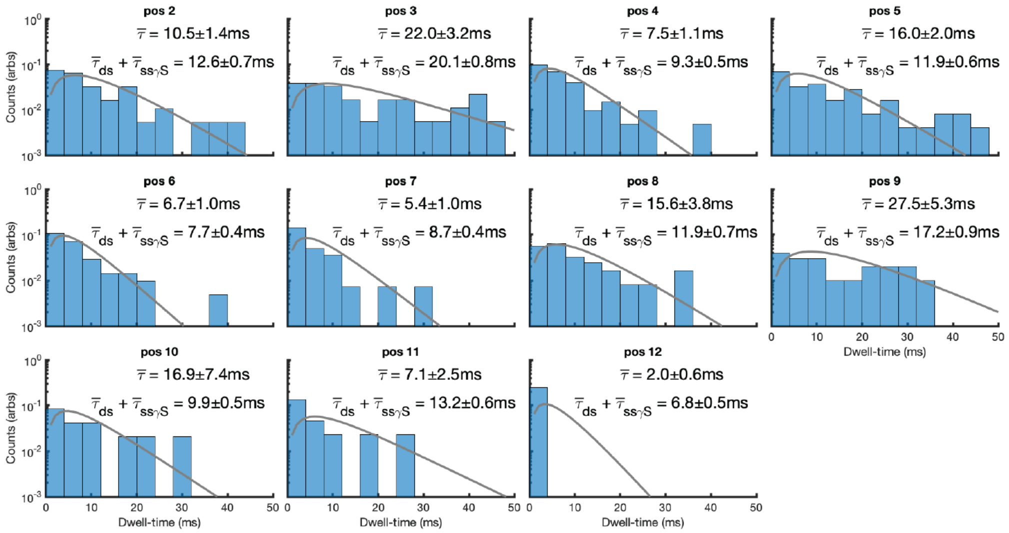

Probability distributions for the durations of individual levels with 600μM ATPγS-only. The y-axis is logarithmic. The histograms were constructed by taking all measurements of the dwell-times of a given level at one 1000 μM ATP condition. The red line is the best fit exponential to the data. Most positions are well described by a single exponential. The mean dwell-time ± S.E.M. are displayed with each distribution. The dwell-time varies from level to level in a statistically significant way, possibly due to interactions between nsp13 and the underlying DNA sequence. Within each subpanel, the mean dwell-time, $\bar{\tau}$, is shown. In our model of ATPγS inhibition, ATPγS-hydrolysis becomes the rate-limiting step during ssDNA translocation and this step precedes the unwinding step. However, in most positions, the duplex-unwinding step is still somewhat slower than ATPγS-hydrolysis though they are a similar order-of-magnitude. This means both processes should contribute to the observed dwell-times. We can use our model of nsp13 motion in combination with our measurements of duplex unwinding at saturating [ATP] and of ATPγS inhibition with ssDNA to make a prediction for the dwell time distributions at each position. This prediction is represented by the gray line in each panel and is given by the convolution of two exponential distributions in which one rate-constant is the mean dwell-time for dsDNA unwinding in the presence of 1000μM [ATP], $\bar{\tau}_{ds}$, and the other rate constant is the mean step dwell time for ssDNA translocation with 600μM ATPγS-only, $\bar{\tau}_{ss\gamma S}$. The mean dwell-time for this distribution, $\bar{\tau}_{ds} + \bar{\tau}_{ss\gamma S}$, is in agreement with $\bar{\tau}$ for each position.

**Figure S21: Comparing forces applied in optical tweezer vs SPRNT experiments**

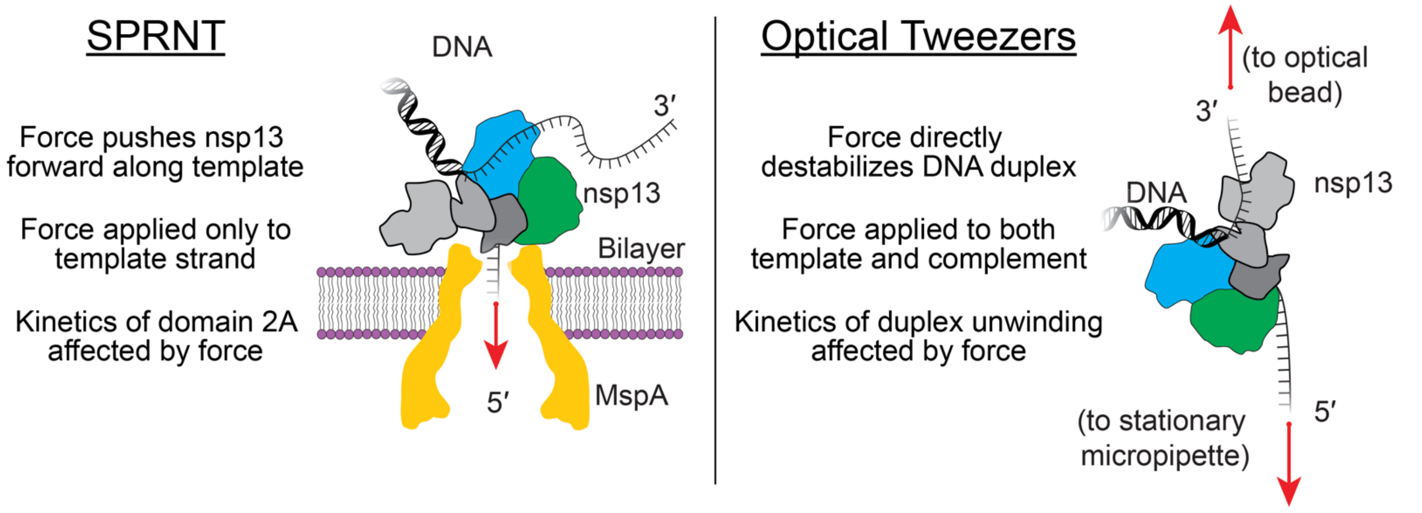

1. Butler, T. Z., Pavlenok, M., Derrington, I. M., Niederweis, M. & Gundlach, J. H. Single-molecule DNA detection with an engineered MspA protein nanopore. *Proc. Natl. Acad. Sci.* **105**, 20647–20652 (2008).
2. Mickolajczyk, K. J. *et al.* Force-dependent stimulation of RNA unwinding by SARS-CoV-2 nsp13 helicase. *Biophys. J.* **120**, 1020–1030 (2021).
3. Derrington, I. M. *et al.* Nanopore DNA sequencing with MspA. *Proc. Natl. Acad. Sci.* **107**, 16060–16065 (2010).
4. Laszlo, A. H. *et al.* Decoding long nanopore sequencing reads of natural DNA. *Nat. Biotechnol.* **32**, 829 (2014).
5. Durbin, R., Eddy, S. R., Krogh, A. & Mitchison, G. *Biological sequence analysis: probabilistic models of proteins and nucleic acids*. (Cambridge university press, 1998).
6. Craig, J. M. *et al.* Revealing dynamics of helicase translocation on single-stranded DNA using high-resolution nanopore tweezers. *Proc. Natl. Acad. Sci.* **114**, 11932–11937 (2017).
7. Laszlo, A. H. *et al.* Sequence-dependent mechanochemical coupling of helicase translocation and unwinding at single-nucleotide resolution. *Proc. Natl. Acad. Sci.* **119**, e2202489119 (2022).
